# Disabling Gβγ SNARE interaction in transgenic mice disrupts GPCR-mediated presynaptic inhibition leading to physiological and behavioral phenotypes

**DOI:** 10.1101/280347

**Authors:** Zack Zurawski, Analisa D. Thompson Gray, Lillian J. Brady, Brian Page, Emily Church, Nicholas A. Harris, Michael R. Dohn, Yun Young Yim, Karren Hyde, Douglas P. Mortlock, Danny G. Winder, Simon Alford, Carrie K. Jones, Heidi E. Hamm

## Abstract

G_i/o_-coupled G-protein coupled receptors modulate neurotransmission presynaptically through inhibition of exocytosis. Release of Gβγ subunits decreases the activity of voltage-gated calcium channels (VGCC), decreasing excitability. A less understood Gβγ–mediated mechanism downstream of calcium entry is the binding of Gβγ to SNARE complexes. Here, we create a mouse partially deficient in this interaction. SNAP25Δ3 homozygote animals are developmentally normalbut impaired gait and supraspinal nociception. They also have elevated stress-induced hyperthermia and impaired inhibitory postsynaptic responses to α_2A_-AR, but normal inhibitory postsynaptic responses to G_i/o_-coupled GABA_B_ receptor activation. SNAP25Δ3 homozygotes have deficits in inhibition of hippocampal postsynaptic responses to 5 HT_1b_ agonists that affect hippocampal learning. These data suggest that G_i/o_-coupled GPCR inhibition of exocytosis through the Gβγ-SNARE interaction is a crucial component of numerous physiological and behavioral processes.

## INTRODUCTION

G protein coupled receptors (GPCR) play a primary role in regulating every physiological function and every organ system. They modulate secretion of hormones throughout the body. In the brain, GPCRs activated by neurotransmitters in turn modulate neurotransmission- and voltage-gated ion channels and are critical to proper functioning of brain circuits as well as their alterations in development, especially learning and memory. GPCR modulation occurs through many mechanisms, including second messengers and phosphorylation^1^. In particular, G_i/o_ GPCRs are inhibitory to secretion throughout the body from the periphery to the central nervous system via direct membrane-delimited action through Gβγ. In neurons presynaptic modulation of neurotransmitter release occurs at every synapse and is important in avoiding overstimulation by autoreceptors that inhibit the release of specific neurotransmitters as well as for normal functioning of brain circuitry by heteroreceptors. G_i/o_ GPCRs inhibit exocytosis presynaptically through three main membrane-delimited mechanisms: by voltage-dependently inhibiting calcium entry through Gβγ binding to voltage-gated calcium channels (VGCC)^2-10^, by Gβγ activation of G protein-coupled inwardly-rectifying potassium channels ^11-13^, leading to membrane hyperpolarization, and by direct interaction of Gβγ with the exocytotic apparatus causing inhibition of exocytosis^14-29^.

An accepted mechanism of presynaptic inhibition involves G_i/o_-coupled GPCR modulation of calcium entry at the active zone^10,30-33^. Free cytosolic Ca^2+^ increases neurotransmitter release with a 4^th^ power non-linear relationship^34,35^ between concentration and exocytosis, and inhibition of Ca^2+^ entry inhibits exocytosis. Direct Gβγ modulation of VGCC has been demonstrated at the presynaptic terminal with calcium-sensitive dyes^25^ and more directly, electrophysiologically at cell bodies and at the calyceal synapse of Held^36,37^. G_i/o_-coupled GPCRs also inhibit the frequency of spontaneous firing and Ca^2+^- independent neurotransmitter release through less well-understood mechanisms^38-44^. Direct inhibition of exocytotic fusion controls exocytosis linearly and does not require changes in diffusible cytoplasmic calcium. Gβγ modulates exocytotic fusion by binding the membrane-proximal C-terminal end of the soluble <u>NSF</u> attachment protein receptor (SNARE) complex that brings the vesicle close to the plasma membrane. We have shown that Botulinum Toxin Type-A (BoNT/A) functionally uncouples this direct Gβγ–SNARE interaction by cleaving the C-terminal 9aa from SNAP25; in addition, a peptide from this region can itself block the presynaptic inhibition^17^.

In biochemical support of the modulation at the exocytotic machinery, we have shown that Gβγ binds directly to ternary SNARE complexes, as well as to the target membrane-associated (t)-SNARE and the individual SNARE components SNAP25, syntaxin1A, and synaptobrevin2/VAMP2, and competes with the binding of synaptotagmin I to t-SNARE and ternary SNARE complexes^16,21^. In addition, we have shown that Gβγ mediates a profound inhibition of exocytosis upon the activation of presynaptic G_i/o_-coupled 5HT_1B_- like receptors^15,16,21,25^.

Modulatory effects of presynaptic GPCRs “downstream of calcium entry” are difficult to measure directly in the presynaptic terminal. This is due to preclusion of direct recording by the small size of the terminal. Many critical questions remain unanswered about the importance of Gβγ regulation of the SNARE machinery in neuromodulation. For example, it is not known which G_i/o_-GPCRs work by this mechanism. In particular, the relative importance of direct inhibition of release at the exocytotic apparatus versus other G_i/o_-GPCR evoked molecular events is unknown. Moreover, it is not known whether multiple Gβγ– dependent mechanisms operate synergistically.

To investigate the role of the Gβγ-SNARE interaction in normal physiology, we created a transgenic mouse deficient in this interaction using the CRISPR/Cas9 strategy. We focused on the SNAP25 target of Gβγ, because of the difficulty of removing Gβγ, as there are 5 Gβ’s and 12 Gγ’s, and because we do not know which of them is involved in interaction with the SNARE complex. However, we know that Gβγ binds the C-terminal of SNAP25, and we have determined that SNAP25Δ3 has a twofold reduction in Gβγ binding and a two-fold reduction in the serotonin 1B receptor (5-HT1BR) mediated inhibition of exocytosis, while evoked release is normal^26,45^. In the current study, we confirmed that C-terminally truncated t-SNARE complexes have a decreased ability to bind Gβγ and inhibit synaptotagmin I-mediated liposome fusion *in vitro*. Furthermore, we created a transgenic mouse with the SNAP25Δ3 mutation. *In vivo* studies reveal that the SNAP25Δ3 mutant animals have significant behavioral and physiological defects in nociception and stress handling, motor coordination, affective behaviors, and cognitive behaviors linked to the central nervous system. This is likely due to suppression of G_i/o_-coupled GPCR-mediated inhibition of exocytosis.

## RESULTS

### Creation of the SNAP25Δ3 mouse via CRISPR-Cas9

To create a mouse deficient in the Gβγ-SNARE interaction, we introduced the SNAP25Δ3 mutation into the eighth exon of SNAP25 via the CRISPR/Cas9 interaction. We inserted the SNAP25Δ3 allele into the wildtype (WT) SNAP25 locus using CRISPR-Cas9 technology. First, sgRNAs targeting the C-terminus of SNAP25 were ligated into the Cas9-containing px330 vector (**Fig. 1A**). Single-stranded homology donors contained the G204* mutation within the C-terminus of SNAP25 and a HindIII restriction site for genotyping, along with appropriate regions of homology to induce homology-driven repair (**Fig. 1A**). These DNAs were co-transfected into B6D2 mouse embryos. Transfected embryos were implanted into dams and pups were born. Heterozygote pups were bred to obtain viable and fertile homozygotes. PCR genotyping of the pups revealed three heterozygous mice that had undergone homology-directed repair out of 32 pups born (**Fig. 1B**). To verify the integrity of the regions surrounding the G204* mutation, we performed Sanger sequencing on PCR products from the eighth exon of SNAP25 (data not shown). No mutations other than the G204* mutation and subsequent restriction site in the 3’ UTR for genotyping were observed. Heterozygous mice were fertile and were bred to C57BL6/J WT mice to yield homozygous offspring.

**Figure 1.**
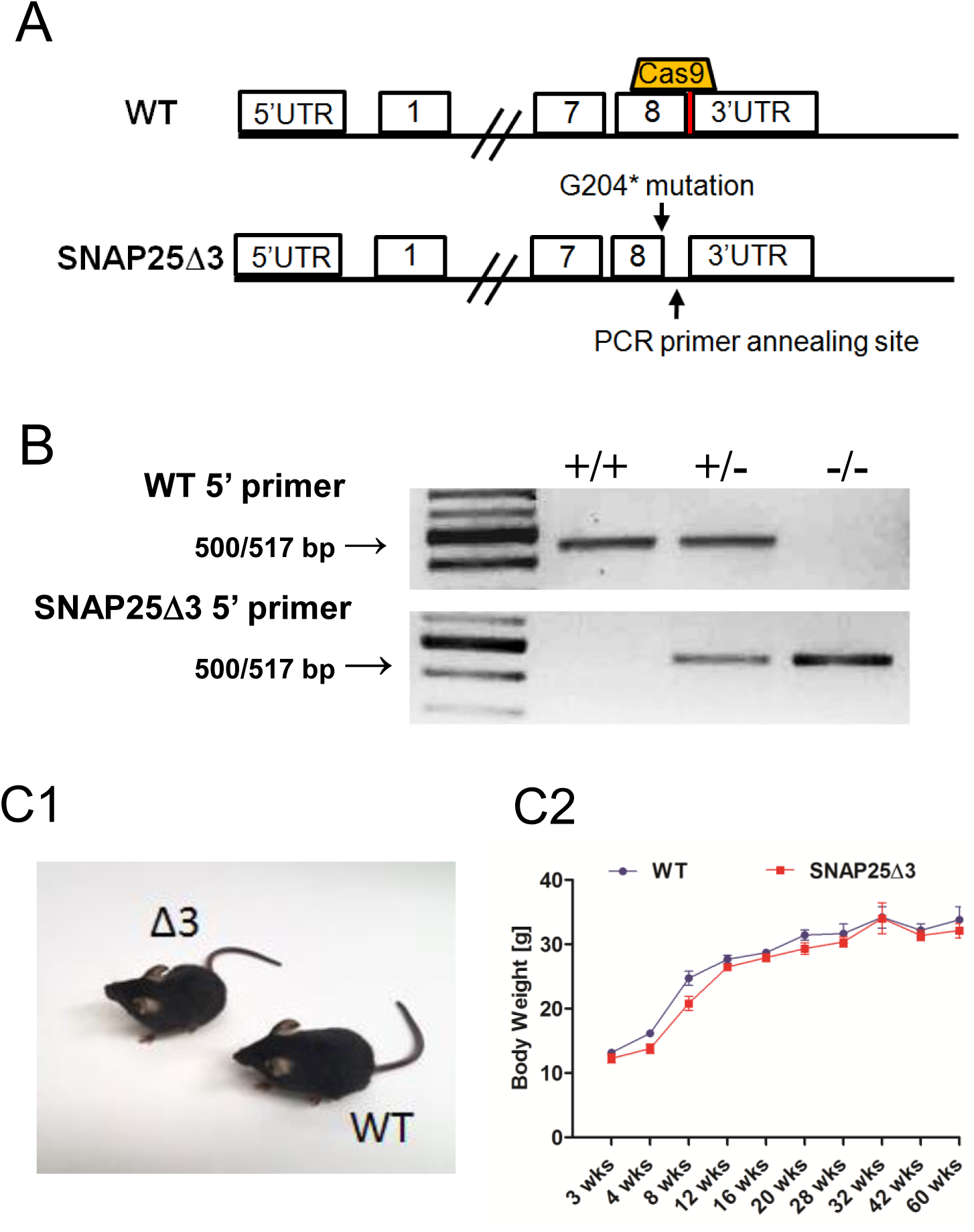
Creation of the SNAP25Δ3 mouse via CRISPR-Cas9. Graphical representation of region on mouse chromosome 2 targeted by the sgRNA (upper panel) cloned into px330 and the subsequent region after homology-directed repair (lower panel) containing the G204* mutation and cloning site. **B**. Agarose gel electrophoresis of PCR products generated from reactions containing two different 5’ primers utilized to genotype WT, heterozygote, and homozygote SNAP25Δ3 littermate animals. The WT 5’ primer corresponds to the WT region on mouse chromosome 2, whereas the SNAP25Δ3 primer corresponds to the region containing the G204* mutation. **C1**. Gross morphology of WT and SNAP25Δ3 homozygotes and (**C2**) growth curves showing the increase in body mass in WT (n= 12-18) and SNAP25Δ3 (n= 12-21) homozygotes over the first 60 weeks of life of several noncontinuous cohorts. **D**. Immunoblot analysis of presynaptic proteins expressed within synaptosomal^100,94,101^ fractions (all except for syt I and VII, and VAMP2) or whole mouse brain lysate (syt I and VII, and VAMP2). A decrease in cysteine string protein (CSP) was found in the presynaptic fraction of SNAP25Δ3 synaptosomes (P<0.03, N=12). No statistically significant difference was found in other proteins. **E**. Immunofluorescence imaging of GABAergic (VGAT) and glutamatergic (vGlut2) immunoreactive appositions^46^ within hippocampal slices taken from adult WT or SNAP25Δ3 homozygotes (n= 4). Anti-HuC/D was utilized asa pan-neuronal marker (Hu proteins are mammalian embryonic lethal abnormal visual system (ELAV)- like neuronal RNA-binding proteins).

Homozygous offspring containing the SNAP25Δ3 mutation were viable at all developmental ages and the general health, appearance, and breeding of these mutant mice was unremarkable relative to their age-matched littermate WT mice **(Fig. 1C1)**. SNAP25Δ3 mutant mice also displayed normal growth curves relative to littermate WT mice, with comparable body weights from 3 to 60 weeks of age **(Fig. 1C2)**. The expression levels of a panel of synaptic proteins present in SNAP25Δ3 mouse brain homogenate were not significantly different from expression levels for these proteins in the WT animals, except for decreased expression of cysteine string protein (CSP) and complexin 1/2, as assessed by immunoblot **(Fig. 1D)**.

Gβγ interactions at presynaptic terminals are ubiquitous and loss of a site of interaction at the presynaptic might cause neurodevelopmental defects. Thus, we utilized immunofluorescence microscopy to investigate differences in neuronal morphology between adult WT and SNAP25Δ3 animals. Neuronal cell bodies were visualized utilizing a mouse primary antibody against HuC/D^46^. Excitatory inputs onto observed soma were visualized using a primary antibody against vGlut2, a marker of glutamatergic synapses, whereas inhibitory inputs were visualized with a primary antibody against vGAT, a marker of GABAergic synapses. No significant differences were detected in the number of synaptic contacts or excitatory/inhibitory inputs between WT and SNAP25Δ3 littermates, supporting the notion that observed phenotypes are due to changes in neuronal signaling rather than changes in synaptic proteins or other gross morphological changes **(Fig. 1E)**.

### SNAP25Δ3 impairs Gβγ competition with synaptotagmin I and inhibition of calcium-Synaptotagmin I-mediated liposome fusion

Gβγ has been shown to mediate its effects on exocytosis via competition with the Ca^2+^ sensor synaptotagmin I (sytI) for interaction for the C-terminal region of SNAP25^16,21^. To investigate the *in vitro* phenotype of the SNAP25Δ3 mutation, we utilized TIRF microscopy to determine whether recombinant Alexa Fluor (AF) 488-labeled sytI C2AB domains bound membrane-associated WT and SNAP25Δ3 t-SNARE complexes differently in lipid membranes, and, additionally, the potency for which purified bovine Gβ_1_γ_1_ could compete with AF-sytI for binding sites on these mutant t-SNAREs. Liposomes consisting of 55% PC/15%PE/29%PS in addition to 1% DiD as a tracer harboring WT or mutant t-SNARE complexes were fused into a lipid bilayer via stimulation with 5mM CaCl_2_. After a superfusate KCl 150 mM and HEPES 5mM wash, AF-sytI with 100 µM free Ca^2+^ was applied in solution over the cover-slip-supported lipid bilayer. Increasing concentrations of Gβγ were then applied in this 100µM Ca2+ solution and AF-sytI fluorescence was measured. Gβγ competed with AF-sytI for binding sites on WT t-SNARE, causing a concentration-dependent reduction in both absolute fluorescence and anisotropy of the AF-sytI signal, with a half maximal effect at 502 ± 151 nM (**Fig. 2A**). Substitution of SNAP25Δ3 into the t-SNARE in the lipid bilayer reduced the maximal effect of Gβγ on AF-sytI anisotropy to 47 ± 13 % of the WT effect (**Fig. 2A**). These data suggest that C-terminally truncated t-SNARE complexes have a decreased ability to bind Gβγ.

**Figure 2.**
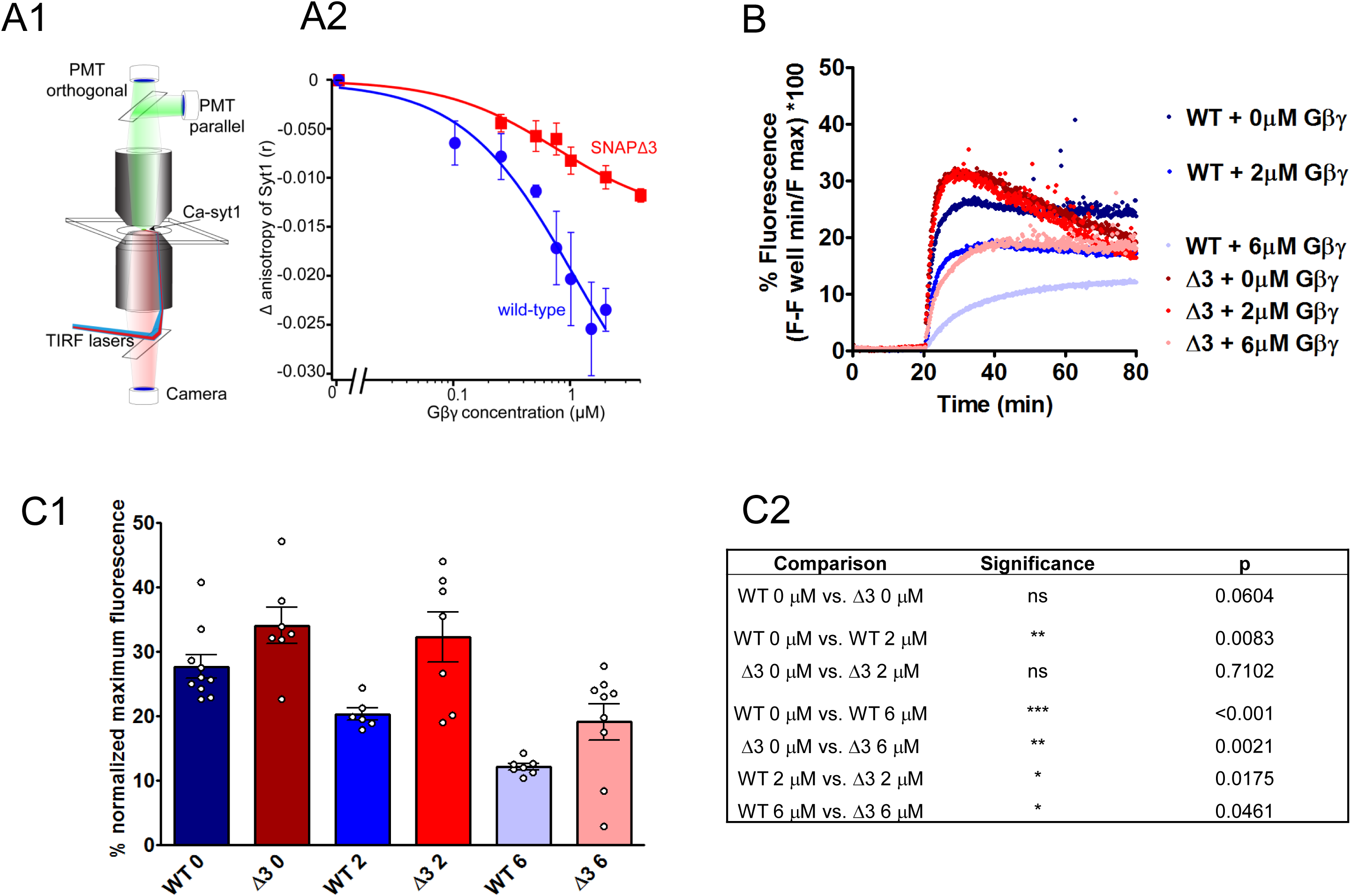
SNAP25Δ3 impairs Gβγ competition with synaptotagmin I and inhibition of calcium-synaptotagmin I-mediated liposome fusion. A. Syt1 competition with Gβγ at t-SNARE complexes in lipid bilayers. (A1) Schematic of the imaging system. A lipid bilayer consisting of 55% PC/15%PE/29%PS /1% DiD harboring t-SNARE complexes was fused to a glass coverslip and imaged using TIRF illumination from a 1.45 NA 60x lens through a laser TIRF illuminator. 1 µM Ca-AF-syt1 was applied over the bilayer (100 µM Ca^2+^). (A2) Graph shows Gβ_1_γ_1_ concentration dependence of tthe change in anisotropy produced by AF-syt I binding to WT (blue) or SNAP25Δ3 (red)-containing t-SNAREs embedded in the lipid membranes. The ability of Gβ_1_γ_1_ to displace AF-syt1 from SNAP25Δ3 t-SNAREs was reduced to 47±13% of its displacement of AF-syt1 from WT SNAP-25, as measured by change in anisotropy(n = 5, p = 0.019). AF-sytI displacement from WT t-SNAREs had an IC_50_ of 502 nM (95% CI: 150nM). **B**. Traces of lipid mixing experiments in which liposomes containing t-SNAREcomplexes made with SNAP25WT or SNAP25Δ3 are incubated with liposomes containing VAMP2 and a FRET pair of NBD-PE and rhodamine-PE in addition to 10μM sytI and Gβ_1_γ_1_. At t= 20 min, 1mM CaCl_2_ is added. **C1**. Bar graph of maximum fluorescence values: 2μM Gβγ significantly inhibits lipid mixing with liposomes containing t-SNAREs made with SNAP25WT (p=0.0083) but not SNAP25Δ3 (p = 0.71), while 6μM Gβγ inhibits significantly less in SNAP25Δ3 liposomes than SNAP25WT (p = 0.0461). **C2**. Table of significance values for lipid mixing experiments (Student’s two-tailed t-test). Experiments were repeated 6-8 times for 6-10 technical replicates.

Gβγ has been shown to inhibit sytI and SNARE-driven lipid mixing assays in a concentration-dependent manner^27^. A more extensive C-terminal mutation of SNAP25, SNAP25Δ9, in which nine residues were truncated had previously been shown to display reduced kinetics and reduced maximum extent of lipid mixing compared to WT SNAP25 in lipid mixing assays^47^. Thus, we investigated whether the SNAP25Δ3 mutant behaved differently from WT controls in reconstituted vesicle fusion studies with SNARE complexes reconstituted into liposomes containing anionic phospholipids^47,48^ (**Fig. 2B**) No significant difference was observed between SNAP25Δ3 and WT SNAP25 with regards to the maximum extent of lipid mixing (**Fig. 2C1**). The basal rate of lipid mixing in the absence of sytI C2AB was not significantly different between SNAP25Δ3 and WT.

The reduced efficacy for Gβγ to displace calcium-bound syt1 at SNARE complexes containing SNAP25Δ3 compared to WT SNAP25 implies that the mutation will modify the ability of Gβγ to interfere with vesicle fusion. Indeed, Gβ_1_γ_1_ was able to significantly inhibit lipid mixing at a concentration of 2μM in liposomes containing t-SNAREs with WT SNAP25, but not SNAP25Δ3 (**Fig. 2C**). Gβ_1_γ_1_ was able to inhibit lipid mixing significantly less well at a concentration of 6μM in liposomes containing t-SNAREs with SNAP25Δ3 than liposomes containing SNAP25 WT. Pairwise comparisons between WT and the SNAP25Δ3 mutant t-SNARE can be found in **Fig. 2C2**. Together, these data suggest that the SNAP25Δ3 mutation impairs Gβγ binding while showing very little effect on lipid mixing or sytI C2AB function, in accordance with previous results^26^.

### SNAP25Δ3 mice exhibited modest alterations in autonomic and somatomotor nervous system functions but enhanced insulin action

Membrane delimited G protein mediated effects are ubiquitous at secretory cells and at presynaptic terminals. Gβγ modifies hormone secretion^24^, and synaptic transmission in the peripheral and central nervous systems^5,6,15,19,28,49^. However, these effects are mediated by more than one target of Gβγ including the SNARE complex. Thus, we next systematically characterized potential changes in the behavioral and/or physiological phenotypes of the SNAP25Δ3 homozygotes. To identify any gross neurological deficits, we performed a modified Irwin Neurological Battery^50^ on male homozygote SNAP25Δ3 and their age-matched littermate WT mice. This test battery evaluated changes in 25 different autonomic and/or somatomotor nervous system endpoints (see **Supplemental Figure 1** for the complete modified Irwin battery results). While the SNAP25Δ3 mice displayed no change in core body temperature, modest alterations in the aggregate Irwin scores for both the autonomic and somatomotor nervous system functions were noted relative to the WT mice reflecting differences in several endpoints, including the corneal and pinna reflexes, leg weakness, and placing loss, a parameter in which the animals is unable to immediately replace its hindlimb in its normal position when moved out of position **(Fig. S1A1** and **S1A2**). These deficits hinted towards the presence of potential impairments in G_i/o_-coupled GPCR signaling and the need to follow-up with more extensive behavioral and physiological characterization of these mice, including assessment of specific autonomic nervous system functions, locomotor coordination, and nociception.

### SNAP25Δ3 mice have impaired motor coordination and altered gait

G protein coupled receptors, including presynaptic receptors, mediate profound effects on locomotor behaviors throughout the neuraxis. These effects range from supraspinal modulation of initiation of behaviors^51^ to modification of output generated within the spinal cord^52^ and involve a number of presynaptically expressed G_i/o_-coupled receptors. To assess potential alterations in locomotor activity in the SNAP25Δ3 mouse under baseline or challenge conditions, we evaluated overall locomotor activity using an open-field test. No significant genotype-based differences in the total distance traveled or rearing behaviors were observed at any interval over the entire 60 min test period in either light or dark conditions for WT and homozygote SNAP25Δ3 mice **(Figure 3A1, 3A2 and Supplemental Fig. S2)**.

**Figure 3.**
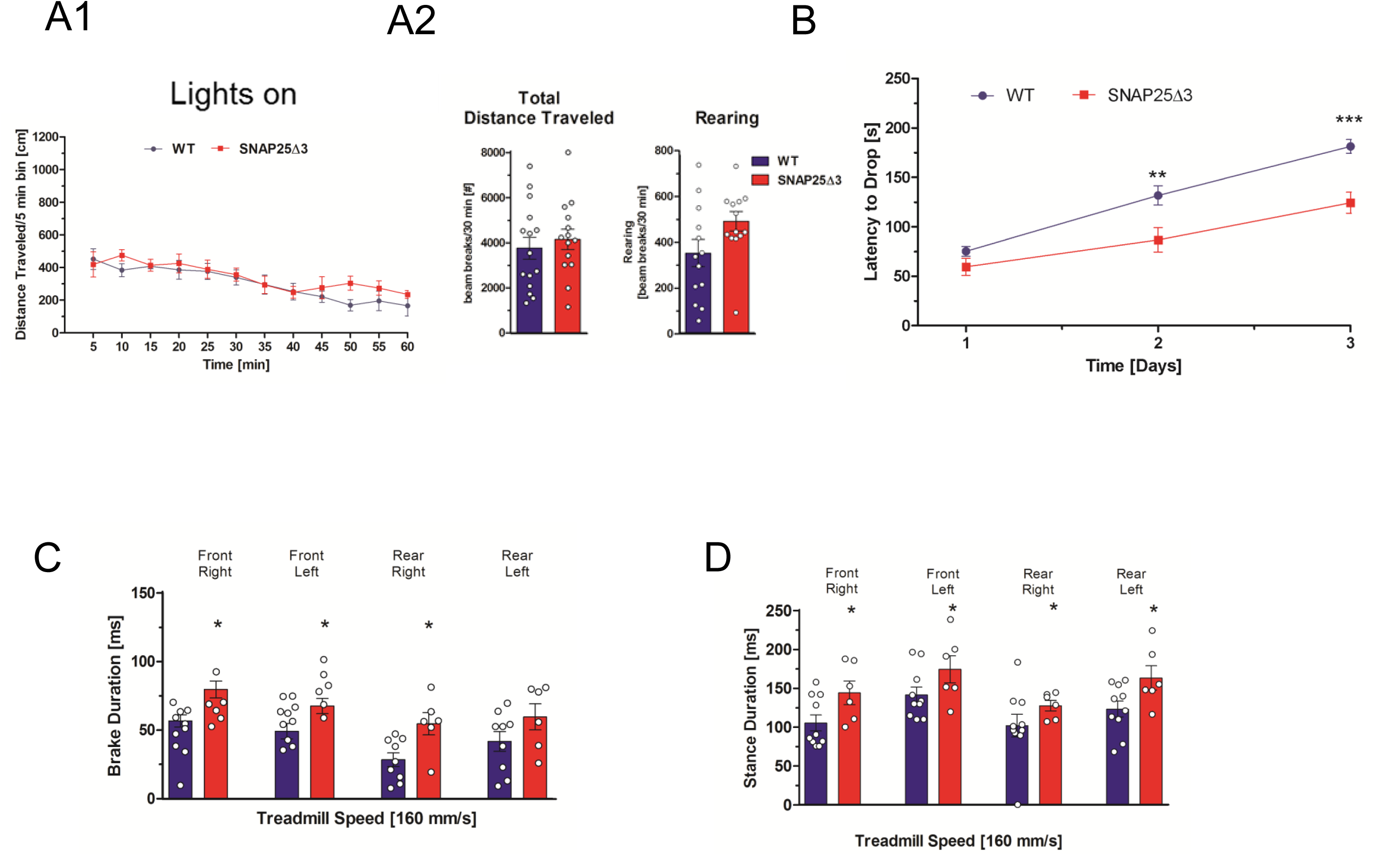
SNAP25Δ3 mutant mice have impaired motor coordination and altered gait. **A1**. Mutant mice have normal locomotor behavior in the open chamber. Plot of distance traveled in five-minute intervals in a brightly illuminated open field for age-matched littermate SNAP25Δ3 homozygotes (red line) (n=14) and WT (blue line) (n= 15) 15 weeks of age. **A2**. Total distance traveled (p=0.8088) and number of rearing movements (p=0.0796) made are plotted below for each genotype. **B**. Plot of latency to drop from an accelerating, rotating beam for 16 week old male WT (n=15) and homozygotes SNAP25Δ3 (n=14) in the rotarod paradigm. Animals were tested daily for three consecutive days. Age-matched littermate SNAP25Δ3 homozygotes had a significantly reduced latency to drop on the second and third day of testing compared to wild-type controls (**, p<0.01, ***, p<0.001, Student’s two-tailed t-test). **C**. Brake duration for each paw as measured by TreadScan^®^ gait analysis in which the paws of a running male age-matched WT or SNAP25Δ3 homozygote are imaged by a camera located beneath a transparent treadmill moving at a speed of 160 mm/s. SNAP25Δ3 homozygotes had significantly greater brake duration than wild-type littermates in each paw (*p < 0.05) except for the rear left paw (p = 0.94,) **D**. Stance duration for each paw as measured by TreadScan^®^ gait analysis in which the paws of a running age-matched WT or SNAP25Δ3 homozygote were imaged by a camera located beneath a transparent treadmill moving at a speed of 160 mm/s. SNAP25Δ3 homozygotes had significantly greater stance duration than wild-type littermates in each paw (*p <0.05).

We evaluated the motor coordination, balance, and learning of the SNAP25Δ3 mice using a multi-day rotarod paradigm where the latency (in seconds) required for a mouse to drop from a constantly accelerating rotating rotarod was measured over three consecutive days. As shown in **Figure 4B**, both the male WT and SNAP25Δ3 mice showed increased motor learning as denoted by increased latencies to drop from the rotarod on each subsequent day of testing. However, the rate of motor learning was decreased in the SNAP25Δ3 mice relative to the WT mice over the testing period, with significant differences observed on the second and third days of testing.

**Figure 4.**
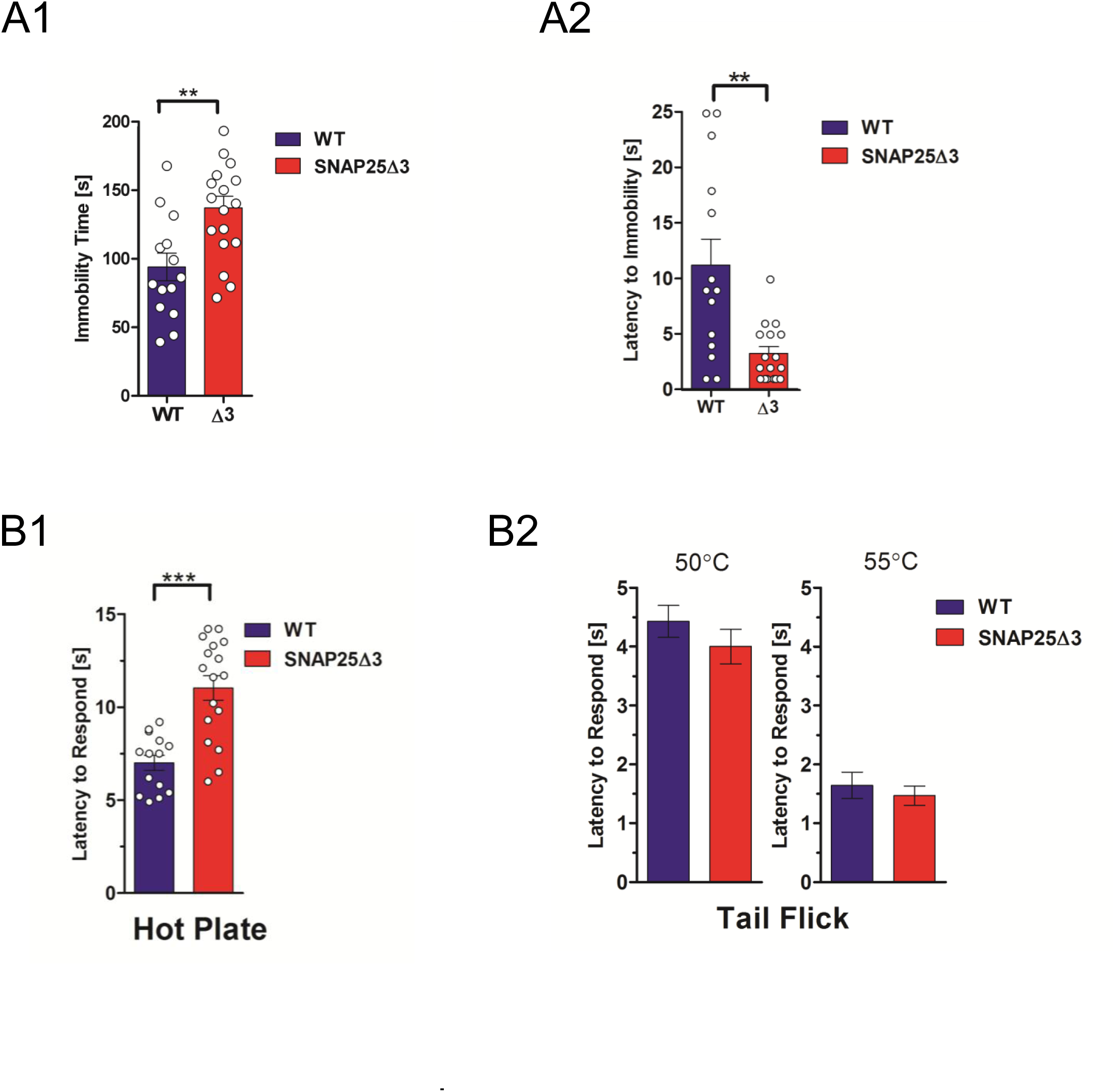
SNAP25Δ3 mutant mice showed altered affect and supraspinal nociception. **A**. SNAP25Δ3 animals show greater immobility in the forced swim paradigm. **A1**. Bar graph shows the time spent immobile subsequent to immersion for 16-17-week old littermate male WT (n= 14) or SNAP25Δ3 (n=17) homozygotes. Immobility time was significantly greater for SNAP25Δ3 than WT littermates (**p< 0.01, Student’s two-tailed t-test) **A2**. Bar graph showing the latency to immobility subsequent to immersion for littermate male WT or SNAP25Δ3 homozygotes in the forced swim paradigm. Latency time before immobility was significantly lower for SNAP25Δ3 than WT littermates (**p< 0.01). **B**. SNAP25Δ3 animals have impaired nociception. **B1**. Bar graph showing latency required for 20 week old littermate male WT (n= 14) or SNAP25Δ3 (n= 17) homozygotes to respond to supraspinal thermal pain in the hot plate paradigm, in which animals are placed on a plate heated to 55°C and the time required to produce a paw movement is measured. Latency time was significantly greater for SNAP25Δ3 than WT littermates (**p< 0.01, Student’s two-tailed t-test). **B2**. Bar graph showing latency required for 21-week old littermate male WT (n= 14) or SNAP25Δ3 homozygotes (n=17) to respond to spinal thermal nociception in in the tail flick paradigm, in which mouse tails are immersed in a hot water bath heated to 50° or 55° C and the time required for tail movement is recorded. No differences were detected between littermate SNAP25Δ3 and WT (p= 0.29 and 0.53 respectively, Student’s two-tailed t-test).

To assess whether the potential differences in the gait of the SNAP25Δ3 mice relative to the WT could have contributed to the differences in rotarod learning over time shown in **Fig 3B**, we evaluated the mice using the TreadScan^®^ gait analysis system^53^ Paw placement and motion of each mouse during running were imaged from below a moving transparent treadmill. Image analysis was performed utilizing TreadScan^®^ software. As shown in **Figure 3C**, SNAP25Δ3 homozygotes exhibited a significant increase in brake duration for all paws other than the rear left paw. Similarly, there was an increase in stance duration in all four paws in SNAP25Δ3 **(Fig. 3D)**. Across other gait parameters, such as gait angle, run speed, track width, stride frequency, stride length, % stance, stride duration, swing duration, and propel duration very minor differences were observed between WT and SNAP25Δ3 homozygotes **(Supplementary Figure S3 A-I)**. Together, these findings suggest the presence of locomotor deficits under challenge paradigms in the SNAP25Δ3 homozygotes, including motor coordination and abnormal gait.

### SNAP25Δ3 mice lack differences in anxiety in the absence of stressors but do have increased rate of helplessness in the forced swim model

Similar to motor control, mood and affect are profoundly modified by GPCRs. An important component of this effect is mediated by presynaptic G_i/o-_coupled receptors including 5-HT_1B_ and α_2A_ adrenergic receptors, cannabinoid CB1 receptors, dopamine D1 and D2 receptors and group II mGluRs. We sought to investigate potential abnormalities in experimental models related to mood and affect in the SNAP25Δ3 homozygote mice. Using the light-dark box paradigm^54^ a preclinical model of anxiogenic-like activity, we assessed whether SNAP25Δ3 mice would spend more or less time in the light portion of the testing chamber in comparison to the age-matched WT littermates. As shown in **Supplementary Fig S4**, there were no significant differences detected between WT and homozygote SNAP25Δ3 mice in the time spent in the light side of the chamber, transitions between the light and dark sides of the chamber, or the distance traveled in the light and dark chambers. These data suggest the absence of an anxiety-like phenotype in SNAP25Δ3 homozygotes in the absence of an external stressor.

We performed the forced swim test^55^, a preclinical model for the evaluation of antidepressant efficacy. Age-matched 16-17 week old WT (n=14) and homozygotes (n=17) were placed into an inescapable, water-filled cylinder and the latency to immobility and the total immobility times were recorded. As shown in **Figure 4**, SNAP25Δ3 homozygotes displayed significantly more immobility time **(Fig. 4A1)** than WTs, as well as a significantly lower latency to immobilization (**Fig. 4A2**). These initial results are consistent with an interpretation of a potential depressive-like phenotype in the SNAP25Δ3 mice.

### SNAP25Δ3 mice show altered supraspinal but not spinal nociception

Due to the potential for presynaptic modulatory effects of GPCRs in pain transmission, we follow-up by investigating whether the SNAP25Δ3 mutation altered either spinal or supraspinal mediated mechanisms of nociception, using the tail flick and hot plate assays, respectively^56^. Surprisingly, SNAP25Δ3 mice exhibited a *decreased* sensitivity to thermal pain in the hot plate paradigm in comparison to age-matched WT littermates **(Fig. 4B1)** as shown by an increased latency to withdraw or lick the footpad after being placed on a 55°C hot plate. However, in the tail flick paradigm (a measure of spinal nociception), no significant change in the latency of a mouse to withdraw its tail from a 55°C hot water bath was observed between age-matched SNAP25Δ3 homozygotes and WT littermates **(Fig. 4B2)**, indicating that SNAP25Δ3 homozygotes have alteration in supraspinal, but not spinal nociception, leading to *increased* pain thresholds.

### SNAP25Δ3 animals have impaired sympathetic responses and loss of α_2a_ adrenergic receptor function

Presynaptic inhibitory effects of GPCRs were first identified in the autonomic nervous system as autoreceptors controlling noradrenaline release^57^. In the Irwin behavioral screening, we noted several changes in autonomic nervous system function in SNAP25Δ3 animals (**Fig S1A1**). One sympathetic system G_i/o_ GPCR that is known to work through the Gβγ-SNARE interaction is the presynaptic inhibitory α_2A_-adrenergic receptor (α_2A_-AR)^19,24^. If the inhibitory effects of α_2A_-ARs are perturbed in the SNAP25Δ3 homozygotes, we might expect to see elevated stress responses, because this receptor is the noradrenergic inhibitory autoreceptor that regulates the amount of norepinephrine (NE) release in the sympathetic system that controls fight and flight responses^58-60^. To investigate whether the SNAP25Δ3 animals have exaggerated stress responses, we performed stress-induced hyperthermia studies in singly-housed age-matched littermate SNAP25Δ3 homozygotes and WT littermate controls^90^. In the stress-induced hyperthermia paradigm^61^, handling-based stress produces an increase in body temperature. Rectal temperatures were recorded before and after manual scruffing with an interval of 15min between each reading. SNAP25Δ3 homozygotes exhibited a twofold greater elevation in body temperature than WT littermates subsequent to handling **(Fig. 5A)**. From this, we concluded that the SNAP25Δ3 mutation produces a stress hypersensitivity phenotype.

**Figure 5.**
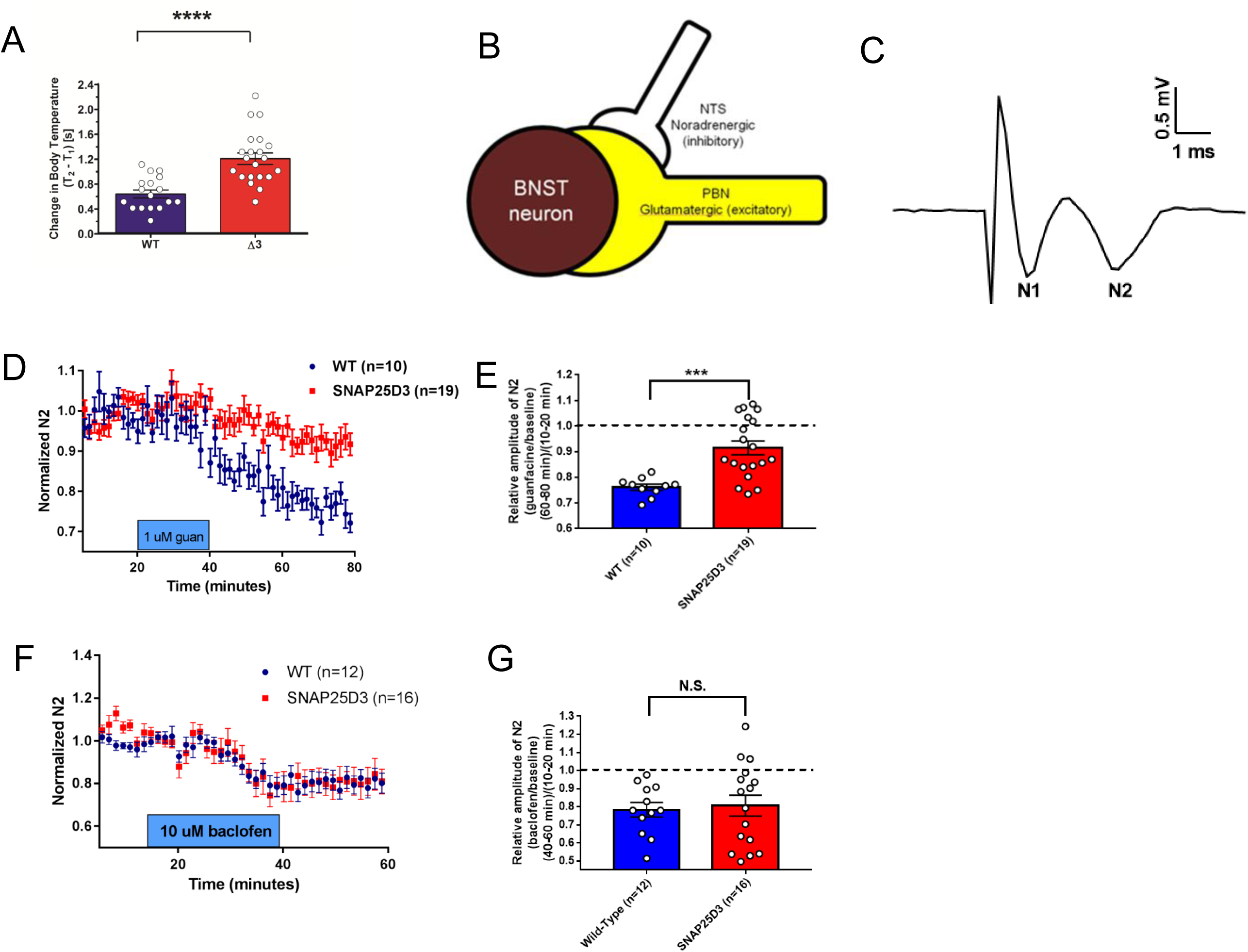
SNAP25Δ3 mice have altered stress responses and impaired α_2a_ heteroreceptor signaling in the BNST. **A**. Bar graph of significant changes in rectal temperature subsequent to handling in singly-housed littermate male WT (n=17) and SNAP25Δ3 homozygotes (n=21) of 13-14 weeks of age (****p<0.00001). **B**. Diagram showing synaptology of α_2a_ heteroreceptor inhibitory signaling on excitatory parabrachial inputs on the bed nuclei of the stria terminalis. **C**. Example field potential from coronal brain slices containing the dorsal BNST illustrating the N1 and N2 downward deflections. **D**. Normalized change in the N2 component of excitatory postsynaptic potentials recorded in the BNST-containing slices taken from WT (in blue) and SNAP25Δ3 homozygote male mice (in red) at an age of >8 weeks. 1uM guanfacine was administered from t= 20 min to t= 40 min. **E**. Bar graph showing relative amplitude of the N2 component of EPSPs at t= 80 min as a fraction of the amplitude prior to the administration of guanfacine at t=10-20min. Guanfacine reduced the N2 component of the EPSP significantly less in slices from SNAP25Δ3 homozygotes than WT (***p < 0.001, Mann-Whitney u-test) **F**. Normalized change in the N2 component of excitatory postsynaptic potentials recorded in the BNST-containing slices taken from WT and SNAP25Δ3 homozygotes at an age of 8-14 weeks. 10uM baclofen was administered from t=20 min to t=40 min. **G**. Bar graph showing relative amplitude of the N2 component of EPSPs at t= 80 min as a fraction of the amplitude prior to the administration of baclofen at t= 10-20min. No significant differences were observed between genotypes(p = 0.92).

Based on our findings of exaggerated stress-induced hyperthermia in the SNAP25Δ3 mutants, we investigated the ability of the α_2A_-AR to inhibit NE release as an autoreceptor^58,62^ as well as glutamate release via heteroceptor mechanisms^63-65^ **(Fig. 5B)** in age-matched littermate WT and mutant animals. To investigate the functions of α_2A_-AR in the SNAP25Δ3 mice, we measured the effect of guanfacine, a partial agonist at the α_2A_-AR, on excitatory glutamatergic transmission in the dorsal bed nucleus of the stria terminalis (dBNST). Guanfacine has been shown to inhibit glutamate release in the BNST via a presynaptic mechanism^64^, at least in part due to inhibition of parabrachial nucleus afferents in the region^65^. Excitatory field potentials were recorded extracellularly as two negative deflections, the tetrodotoxin-sensitive fiber volley potential N1 and the AMPAR antagonist CNQX-sensitive synaptic potential N2^63^ **(Fig. 5C)**. As expected, bath application of guanfacine (1 μM) reduced the amplitude of the N2 in WT animals (WT: 23.9. ± 1.18% of baseline n=10 slices from 7 animals). Bath application of guanfacine (1 μM) attenuated the amplitude of the N2 in SNAP25Δ3 animals (SNAP25Δ3: 8.6 ± 2.70%, n= 19 slices from 10 animals) **(Fig. 5D)**. When compared directly, the reduction in N2 observed in SNAP25Δ3 mice was significantly reduced relative to the change in WT animals **(Fig. 5E)**. To determine whether these effects were specific to the α_2A_-AR or were translatable to other G_i/o_-coupled GPCR, we examined the effects of the GABA_B_ agonist baclofen on excitatory transmission in the BNST. Bath application of 10 μM baclofen inhibited N2 amplitude in both WT (WT: 21.7%, n=12 slices from 4 animals) and SNAP25Δ3 mice (SNAP25Δ3: 19.4%, n = 16 slices from 5 animals) **(Fig. 5F)**. There was no difference in this inhibition when compared across genotypes **(Fig. 5G)**.

#### 5-HT_1b_- and Gβγ–SNARE-mediated inhibition of exocytosis are impaired in homozygous *SNAP25Δ3* mutants, while GABA_B_- and Gβγ–VGCC mediated inhibition of exocytosis is normal

To understand the effect of disruption of the Gβγ-SNARE interaction on electrophysiological responses, we studied the 5HT_1b_-mediated inhibition of hippocampal CA1 to subiculum neurotransmission (**Fig. 6A)**. The subiculum is an essential structure that receives the majority of its input from the hippocampus proper and projects to the entorhinal and other cortices, as well as a variety of subcortical regions. As such, the subiculum plays an important role in spatial and mnemonic information processing, and has been implicated in regulating the response of the hypothalamic-pituitary-adrenal (HPA) axis to stress^66^.

**Figure 6.**
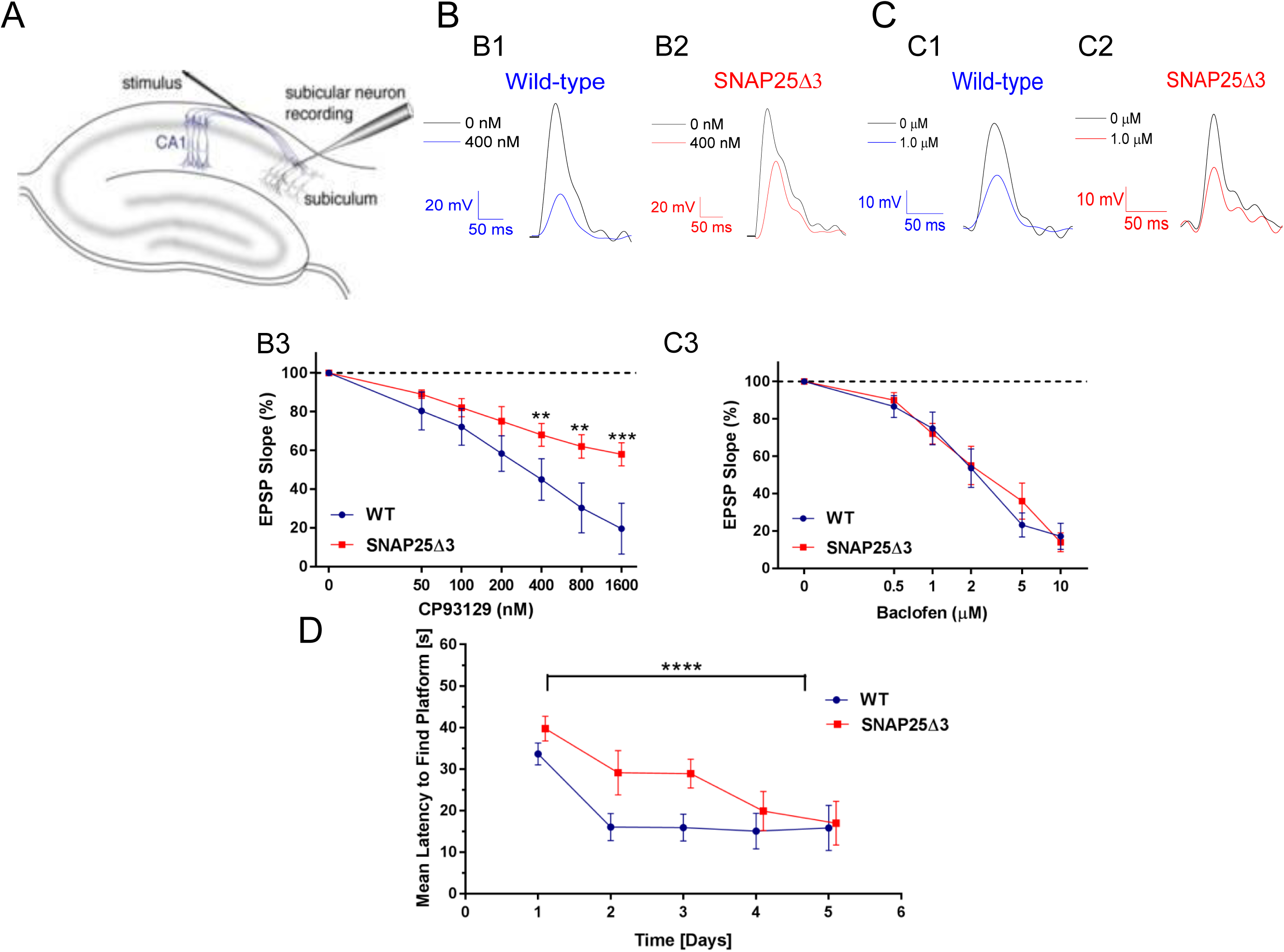
SNAP25Δ3 homozygotes have impaired G_i/o_-coupled GPCR signaling in CA1/subiculuar hippocampal neurons and show impaired hippocampal spatial learning. **A**. Diagram of hippocampal field recording paradigm. Stimulation via bipolar electrodes over the CA1-subicular pathway evoked field EPSPs recorded in basal dendrites of subicular pyramidal neurons in AP5 (50 µM) and bicuculline (5µM) to isolate AMPAR-mediated responses. **B**. Traces from CA1-subicular recordings in WT (**B1**) and SNAP25Δ3 (**B2**) slices at 0 and 400 nM CP93129. **B3**. Dose response of the effect of CP93129 on the AMPA component of these field EPSPs from 6-week old littermate WT (in blue) or SNAP25Δ3 homozygotes (in red). Amplitudes were normalized to the control response. CP93129 was significantly more potent in WT than SNAP25Δ3 (**p = 0.0068, 400 nM; **p = 0.0035, 800 nM; ***p = 0.001, 1600 nM; Student’s t-test). **C**. Traces from field recordings in wild-type (**C1**) and SNAP25Δ3 (**C2**) slices at 0 and 1.0 µM baclofen. **C3**. Dose-response of the effect of baclofen on field EPSPs recorded in the WT or SNAP25Δ3 hippocampal slices. No significant differences were detected by genotype. **D**. Comparison between age-matched littermate WT and SNAP25Δ3 homozygotes in the acquisition of the Morris Water Maze Task over a 5 day trial period by genotype (p<0.05) and time (p<0.0001).

Since 5HT_1B_ receptors have been shown to inhibit synaptic transmission at CA1 to subicular synapses by modulation of the Gβγ-SNARE interaction^25^, we compared the effects of application of the 5HT1BR selective agonist CP93129 on hippocampal slices prepared from age-matched WT or SNAP25Δ3 animals during field potential recordings in the subiculum after stimulation of CA1 pyramidal neuron axons. The GABAAR and NMDAR antagonists (bicuculline, 10 µM; D-2 amino-5-phosphonopentanoate, D-AP5, 50 µM respectively) were applied to allow focus on AMPAR-mediated EPSPs. The slope of excitatory postsynaptic potentials recorded in WT animals (n= 8 slices from 6 animals) was inhibited in a concentration-dependent manner; specifically 400, 800, and 1600 nM of CP93129 caused a significant 50, 70, and 80% decrease in the EPSP slope, respectively (**Fig. 6B1, 6B3**). By contrast, in hippocampal slices from the SNAP25Δ3 animals (n= 5 slices from 5 animals), 400, 800, and 1600 nM of CP93129 caused only 30, 35, and 40% decreases in the EPSP slope, respectively (**Fig. 6B2, 6B3**), a significant difference from EPSP suppression in WT mice. Overall, these data indicate that 5-HT_1b_-mediated inhibition of exocytosis is decreased in the SNAP25Δ3 animals, suggesting that disruption of 5HT_1b_-mediated Gβγ-SNARE interaction in transgenic animals attenuates presynaptic inhibition.

#### No change in GABA_B_–mediated inhibition of neurotransmission from the CA1 hippocampal region to the subiculum is detected in SNAP25Δ3 animals

To examine if synaptic transmission, controlled by Gβγ inhibition of calcium through VGCCs, was altered in SNAP25Δ3 animals, we measured GABA_B_-mediated inhibition of neurotransmission in the hippocampal CA1 to subicular synapse. Comparing the effects of application of the selective GABA_B_ receptor agonist, baclofen, on field potential recordings in WT and SNAP25Δ3 animals, we found that there was no change in GABA_B_-mediated inhibition in the SNAP25Δ3 animals (**Fig. 6C1-3**). These data are consistent with the interpretation that Gβγ regulation of calcium entry by baclofen is normal in the SNAP25Δ3 animals(n= 5 slices from 5 animals for both WT and SNAP25Δ3).

In order to determine whether the SNAP25Δ3 mutation impacted any cognitive functions in the mutant mice, we assessed potential differences between age-matched littermate WT and homozygotes in the Morris water maze task, a well-established preclinical model of hippocampal-mediated learning and memory^67,68^ As shown in **Fig. 6D**, the WT mice quickly learned the position of the hidden platform in this test by day 2 of training. In contrast, homozygous SNAP25Δ3 mice required over 4 days of training to accurately identify the position of the hidden platform, a significant difference in acquisition of this learning/memory.

#### Gβγ regulation of calcium entry and direct Gβγ inhibition of exocytosis at SNARE provide synergistic inhibition of postsynaptic responses

**Fig 7A** shows a schematic of the two distinct Gβγ-mediated neuromodulatory mechanisms, GABA_B_- and Gβγ-mediated inhibition of VGCC and Ca^2+^ entry, and 5-HT_1B_- and Gβγ-mediated inhibition of exocytosis via binding the SNARE complex…Consistently, Gβγ-SNARE inhibition is more dramatic at the first spike of a train and decreases with calcium buildup, while inhibition through VDCCs is consistent throughout the train^69^. Because we have shown that there is a temporal pattern to the effects of Gβγ on SNARE mediated exocytosis^69^ we also examined trains of stimuli. When CA1 pyramidal neuron axons were stimulated repetitively to evoke EPSCs and subicular neurons were whole-cell clamped, the inhibition of the postsynaptic response by the 5-HT1bR selective agonist CP93129 was not sustained, but was attenuated by each consecutive stimulus (blue line, **Fig 7B**). When plotted as a ratio of inhibition of the first to the fifth stimulus, the response was attenuated 4.6±0.8-fold by the fifth stimulus in CP; in baclofen this ratio was 1.1±0.2 (**Fig 7C**). This attenuation in response to CP is interpreted to be due to calcium buildup, allowing sytI to compete more effectively with Gβγ for binding to the SNARE complex^21,70^.

**Figure 7.**
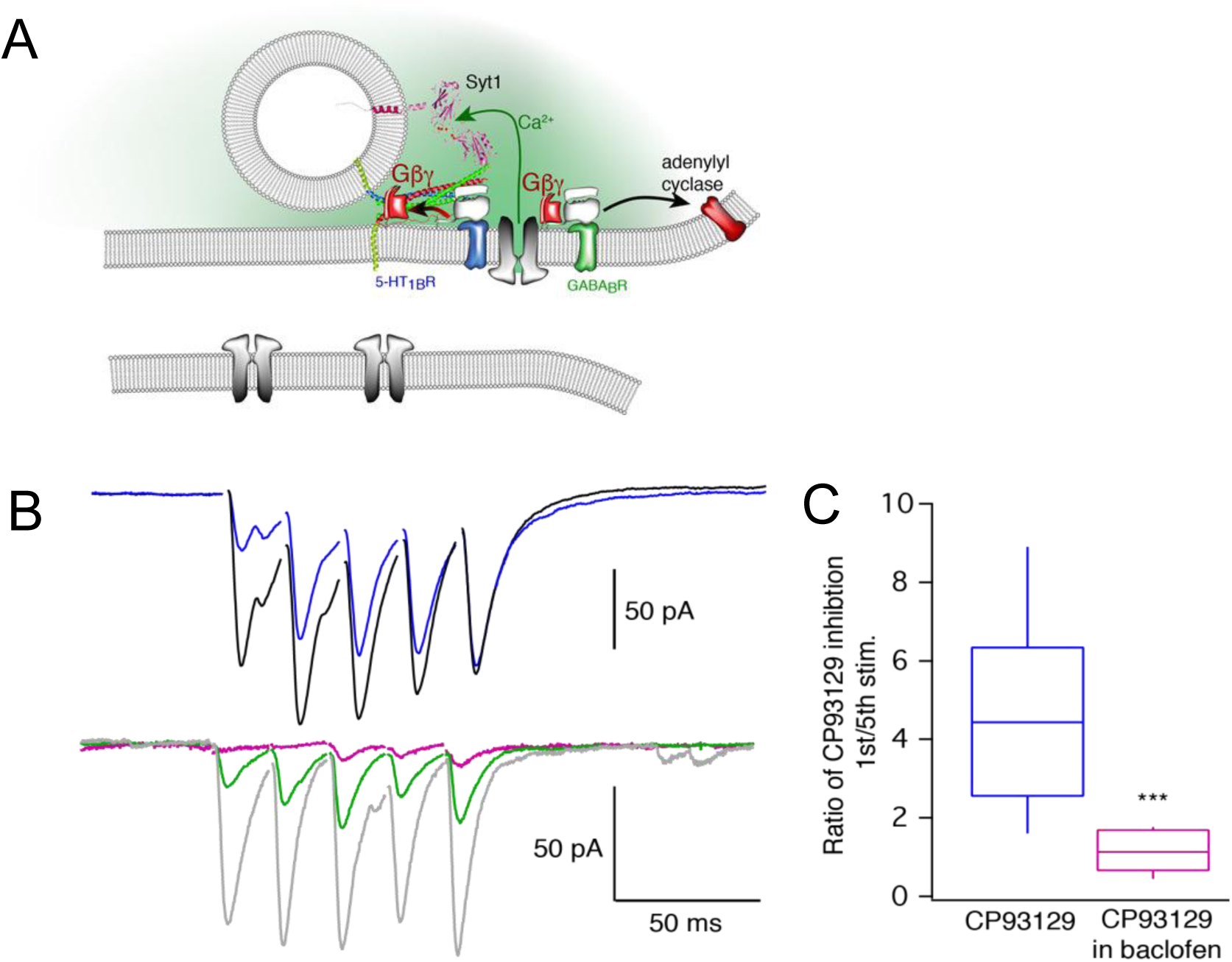
Synergy between 5-HT_1B_ and GABA_B_ at the CA1/subicular synapse. **A**. Schematic of targets within the presynaptic terminal for G_i/o_-coupled GPCRs. In CA1 terminals, 5-HT1bRs Gβγ to bind SNAREs and GABA_B_ receptors release Gβγ to inhibit Ca^2+^ channels. Synergistic effects of 5-HT1bRs and GABA_B_R_s._ **B**. Stimulation of the CA1-subicular pathway evoked whole cell recorded EPSCs in subicular pyramidal neurons. During repetitive stimulation, CP93129 (400 nM, blue) substantially inhibited the first response, but response amplitudes recovered during the stimulus train. The ratio of inhibition of the 1^st^ vs. the 5^th^ response was 4.6±0.8. Baclofen (1 µM, green) uniformly inhibited EPSCs throughout the stimulus train; the ratio was 1.1±0.2. Addition of CP93129 + baclofen, pink) substantially inhibited responses throughout the stimulus train (***p=0.0002). **C**. Quantitation of the effects of CP93129 (400 nM) alone and after addition of baclofen (1 µM).

By contrast, inhibition of calcium entry by baclofen is not phasic in nature, generating a similar inhibition of postsynaptic responses at each stimulus (green line, **Fig 7B**). To determine whether these distinct mechanisms of presynaptic inhibition were additive or more than additive, implying synergism, we examined the impact on inhibition of postsynaptic responses with a simultaneous administration of CP93129 (400nM) and baclofen (1μM). The inhibitory response to this dual stimulation of 5-HT_1b_ and GABA_B_ receptors was much greater than additive (red line, **Fig 7C**). We previously showed that microarchitecture controls GPCRs that work on one or the other mechanism^25^ (**Fig. 7A**), but this is the first time we have shown that these mechanisms can interact presynaptically.

## DISCUSSION

We have created a transgenic mouse that contains a 3aa truncation of SNAP25 at its C-terminus, the SNAP25Δ3 mouse, based on our published evidence that this mutant binds Gβγ less well and Gβγ-mediated modulation of exocytosis is impaired^26^. This mouse model allows us to probe *in vivo* effects of decreasing the neuromodulatory effects of G_i/o_ GPCRs that work by liberation of Gβγ to bind to the SNARE complex and inhibit exocytosis directly. Though this modulatory mechanism has been described *in vitro*^14-17,21,26^, this is the first report of *in vivo* phenotypes caused by its loss.

### The extreme C-terminus of SNAP25 is a conserved region important for Gβγ binding

Multiple independent groups have demonstrated that the SNAP25Δ3 mutant is not deleterious to exocytosis in neurons and chromaffin cells^26,45^. Furthermore, no other SNARE-binding proteins identified to date have been shown to utilize the C-terminal three residues of SNAP25 as a critical site of interaction except Gβγ. This allowed us to selectively examine the impact of Gβγ–SNARE interactions *in* vivo. Gβγ directly competes with sytI for binding to t-SNARE made with SNAP25 WT, while it is less able to compete with sytI for binding to tSNARE made with SNAP25Δ3 (**Fig. 2A)**. We confirmed that t-SNARE complexes made with SNAP25Δ3 do not have defects in synaptotagmin I-stimulated liposome fusion (**Fig. 2C**), with a maximal extent of lipid mixing not different from wild-type t-SNAREs. In addition, Gβγ-mediated inhibition of Ca^2+^-sytI and SNARE-dependent lipid mixing was reduced with t-SNARE complexes made with SNAP25Δ3 (**Fig. 2C1, C2**). Taken together, these data suggest that the specific phenotypes observed in SNAP25Δ3 homozygote neurons are due to a decreased ability of Gβγ to bind SNARE complexes and displace sytI at the terminal.

Gβγ binding to the C-terminal region of SNAP25 was first identified by the effects of BoNT/A cleavage of its C-terminal 9 residues on synaptic modulation^17^. Gβγ liberated from 5HT_1B_ receptors inhibits exocytosis by displacing Ca^2+^-dependent sytI binding to t-SNARE^16,21,27^. This results in a phasic inhibition of synaptic transmission as calcium buildup during a train of action potentials allows sytI to compete more effectively with Gβγ for binding to the SNARE complex^21,70^. As predicted, injection of EGTA was shown to eliminate the phasic inhibition during trains^21,70^. Biochemically, competition between Gβγ and sytI depends upon the concentrations of both Gβγ and sytI, demonstrating this competition between Gβγ and Ca^2+^-sytI interactions with the t-SNARE complex^21^. The ability of Gβγ V displace sytI is lost in SNAP25 mutants lacking identified Gβγ interaction sites, both in models of binding in aqueous solution and in models in which the SNARE complex is embedded in lipid bilayers^16,21,27^. In this study, we have confirmed that Gβγ competes less well with sytI for binding to t-SNARE made with SNAP25Δ3 than with t-SNARE made with SNAP25 WT (**Fig. 2A)**. As predicted, Gβγ-mediated inhibition of Ca^2+^-sytI and SNARE-dependent lipid mixing was reduced with t-SNARE complexes made with SNAP25Δ3 (**Fig. 2 C1, C2**).

Conservation of the amino acid sequence in the C-terminus of SNAP25 is extremely high, particularly with regards to the C-terminal nine residues, which are conserved between mammals, birds, and reptiles, pointing to the evolutionary importance of this region. Vectorial assembly of the SNARE complex proceeds from the N- to the C-terminus^71^, and the C-terminal region of the zippering SNARE complex is crucial for the generation of force leading to fusion pore formation. A truncation of the C-terminal 9 residues of SNAP25 leads to smaller foot currents and reduced fusion pore conductances^72,73^, consistent with impaired vesicle-surface fusion events. Thus, it is very interesting that Gβγ binding to the extreme C-terminus modifies fusion properties^74^ and inhibits exocytosis, suggesting that it may interfere with fusion pore opening. Structural studies on the nature of the Gβγ-SNARE complex would elucidate the nature of this inhibitory mechanism.

The high conservation of this region may also be required for tight control over exocytosis via regulation by G_i/o_-GPCRs of the Gβγ-SNARE interaction, which is present even in extremely primitive vertebrates such as lamprey. In fact, we first discovered this phenomenon in the primitive cartilaginous fish sea lamprey, which had the advantage of giant axons in their spinal cord, because of lack of myelination, allowing us to inject proteins presynaptically^15,17^.

### Clear in vivo delineation of Gβγ functions in regulating exocytosis

In SNAP25Δ3 animals, both the α_2A_-ARs and the 5HT1bR, which have previously been shown to work at least partially through Gβγ-SNARE^19,25,75^ are unable to inhibit exocytosis to the same extent as WT littermates. By contrast, in both the hippocampus and the BNST of SNAP25Δ3 animals, there is no effect on Gβγ inhibition of exocytosis through the GABA_B_ receptor, which is known to work through modulation of calcium entry. This is clear evidence that Gβγ responses through voltage-gated calcium channels- the major other membrane-delimited mechanism regulating neurotransmission-have not been perturbed by the SNAP25Δ3 mutation. Interestingly, receptors working through both modulatory mechanisms can be present in the same synapse, as shown in recent electrophysiological studies in hippocampus^25^.

### Different roles of the two mechanisms of presynaptic neuromodulation

While we do not know in most cases which mechanisms are activated by different GPCRs, and, in fact, this may be different in different neurons^29^, we do know that individual synapses can contain multiple G_i/o_ GPCRs. As previously reported^69^ and as follows from direct Gβγ competition with sytI SNARE interactions (**Fig 2A**), the effects of Gβγ-SNARE interaction are attenuated in response to trains of action potentials, because of buildup of residual Ca^2+^ during the course of a stimulus train^17^. As Ca^2+^ concentrations in the presynaptic terminal rise, calcium-sytI becomes a better competitor of Gβγ at tthe SNARE complex and Gβγ inhibition decreases^21^ **(Fig. 7A**). This opens the possibility that other signal transduction mechanisms within the presynaptic terminal can act synergistically with the Gβγ–SNARE interaction. For example, at the CA1-subicular synapse, 5-HT_1B_ agonists cause Gβγ to interact directly with the SNARE complex. At the same synapse, GABA_B_ receptors inhibit presynaptic Ca^2+^ entry, so during trains GABA_B_ receptor activation reduces action potential-mediated increases in Ca^2+^ concentration. When both receptors are activated, decreased residual Ca^2+^ buildup by GABA_B_ receptor activation substantially magnifies the 5-HT mediated inhibition during the stimulus train, leading to profound presynaptic inhibition. Such a dual modulation, which has never been shown previously, substantially increases the repertoire of presynaptic inhibition, potentially providing much greater control of transmission than either mechanism alone.

The two modulatory mechanisms have very different characteristics. Gβγ’s inhibition of VDCCs is highly sensitive and powerful, but easy to saturate, whereas Gβγ-SNARE interaction right at the final step of membrane fusion is stoichiometric (i.e. one or two Gβγ per SNARE) and phasic, based on Ca2+-syt interactions, and thus may provide a much broader dynamic range of modulation that is more difficult to saturate. SNAP25 represents a highly circumscribed target of Gβγ. Gβγ interacts directly with the SNARE complex providing a mechanism to modulate vesicle fusion, but without other known downstream effectors. The C-terminal region of SNAP25 has been shown to be critical for forces associated with fusion pore, formation and stability^71,73,76^. Indeed, consistent with this important role of the C-terminal region in fusion pore dynamics, Gβγ interaction with this target modifies fusion properties themselves as a mechanism of presynaptic inhibition^70,74,77^. Receptors that simultaneously target presynaptic Ca^2+^ channels, in contrast, mediate a much broader range of presynaptic effects, from modifying action potential shape, to altering release probability, as well as modulating Gβγ effects at the SNARE complex by altering Ca^2+^ accumulation during stimulus trains.

### Dramatic effects of the SNAP25Δ3 mutation on stress

As expected, stress responses in SNAP25Δ3 homozygotes were significantly enhanced, given that the major adrenergic autoreceptor which inhibits norepinephrine release works through this mechanism^78^. This behavioral phenotype was followed up with treatment of brain slices from these mice with guanfacine, a α_2A_-AR selective agonist. We showed that the effect of guanfacine on excitatory glutamatergic transmission in the dorsal BNST, a component of the extended amygdala, was nearly eliminated in homozygous SNAP25Δ3 mice. Previous work has suggested that this receptor presynaptically regulates excitatory input in the extended amygdala via a Gβγ and BoNT sensitive mechanism^19^. Our results here provide further support for such a mechanism at these glutamatergic synapses. Heteroreceptor effects of α_2A_-AR agonists such as those shown here have been shown to have dramatic outcomes on both physiologies within the extended amygdala^63-65^ as well as on whole-organism behavior^78,79^. Specifically, heteroreceptor α_2A_-ARs have been shown to be responsible for the sedation, anesthetic sparing, hypothermia, analgesia, bradycardia, and hypotension induced by systemic administration of α_2A_-AR agonists^56^. This contrasts with the autoreceptor α_2A_-ARs, which are responsible for physiologic feedback inhibition of NE release and spontaneous locomotor activity^78^. The relative contributions and deficits of auto- and hetero-receptor α_2A_-ARs to the stress-related phenotypes observed in the SNAP25Δ3 mouse remain unclear and will be the focus of future studies investigating the mechanisms underlying the ability of α_2A_-AR agonists to block the ability of stress to impact neurotransmission^80^, depression- and anxiety-related phenotypes^81,82^ and drug-seeking behavior in rodent models of addiction^83-85^.

### Chronic disruption of the Gβγ-SNARE interaction leads to a number of behavioral and physiological phenotypes

Prior studies have highlighted the physiological settings in which the Gβγ-SNARE mechanism is critical for the regulation of vesicle release by G_i/o_-coupled GPCRs in a variety of secretory cell types, including multiple populations of neurons in the amgydala, cerebellum, spinal cord, and hippocampus^15,19,20,49^, chromaffin cells^22^, the beta cells of the islets of Langerhans^24^, and cone photoreceptors^28^. Here, we expand upon these existing findings in multiple ways. For many years, presynaptic receptors have been thought to work mainly through modulation of calcium entry by inhibiting VGCC^2-10^, or by activating GIRK channels^86,87^. Using the SNAP25Δ3 mouse as a model of deficiency in the Gβγ-SNARE pathway, we demonstrated that multiple deleterious behavioral phenotypes associated with stress, locomotion, pain processing and spatial learning were detected upon selective inhibition of Gβγ-SNARE interaction, as well as increased immobility in forced swim tests associated with depressive-like phenotype, implying downstream physiological and pathophysiological consequences for disruption of the Gβγ-SNARE pathway.

Though severe physiological consequences exist for SNAP25-null homozygotes, neither neonatal lethality nor gross deficits in exocytosis are observed in the SNAP25Δ3 homozygotes. Physiological consequences for disruption of SNAP25 function are also observed in the SNAP25b-deficient homozygotes, in which the exon specific to SNAP25b is replaced by a second SNAP25a exon^88^, producing spontaneous seizures and a number of behavioral deficits. Locomotor deficiencies in the rotarod test were observed in the SNAP25Δ3 homozygotes, similar in magnitude to the SNAP25 I67T homozygotes termed the “blind-drunk mouse”^89^, although the SNAP25Δ3 homozygotes lacks any of the deficits observed in the light-dark box test present in the I67T homozygotes. It is uncertain if the spatial learning deficits observed in the SNAP25b-deficient mouse in tasks such as the Morris water maze would similarly be observed in the SNAP25Δ3 homozygotes. However, neither deficits in the elevated zero/plus maze paradigm nor spontaneous seizure activity were observed in the SNAP25Δ3 homozygotes. A comparison of the phenotypes of the highly selective mutation in the SNAP25Δ3 mice versus phenotypes detected in the SNAP25 null and SNAP25 I67T mice will help identify the particular molecular interactions, and the affinity/probability of these interactions, for more selective therapeutic target strategies, especially since expression levels of other presynaptic proteins, such as syntaxin 1a and Munc18, were not altered in SNAP25Δ3 homozygotes.

The significant phenotypes displayed by SNAP25Δ3 homozygotes in many behavioral paradigms are clear evidence that G_i/o_ GPCR regulation of Gβγ binding to the exocytotic fusion machinery is required for normal physiological and behavioral function. Based on the profile of these physiologic and behavioral alterations, we are particularly interested in the GPCR identity and circuitry basis of the presynaptic inhibitory action of GPCRs underlying these changes. Unfortunately, this is not possible to determine in these animals, since we do not know which G_i/o_-coupled GPCRs work through this mechanism. In addition, since the SNAP25Δ3 mutation is found in all cells, further circuit-based studies will need to be carried out to understand the basis of the SNAP25Δ3 phenotype. For example, viral expression of the SNAP25-8A mutation that has no affinity for Gβγ^14^ will be used in future studies to further define the circuitry underlying these behavioral phenotypes. Nevertheless, some circuit implications for these receptors can be made from experiments targeting specific receptor subtypes, with known and selective efficacy at the SNARE complex. These effects are consistent with defects found in the SNAP25Δ3 homozygous mice. In the lamprey spinal cord 5-HT_1B/1D_-like receptors are found on glutamatergic terminals of the descending motor command system and the locomotor pattern generator. Activation of these receptors in lamprey mimics the known effects of 5-HT application in all vertebrate models of locomotion, in which locomotor frequency is lowered^70,90^ Loss of these receptor mediated effects in intact animals would be expected to lead to a loss of locomotor coordination and a reduction in the range of locomotor frequencies the animal can achieve.

These studies provide a basis for the hypothesis that the Gβγ-SNARE interaction and modulation of exocytosis downstream of presynaptic calcium entry is an important neuromodulatory mechanism in a large diversity of circuits, mediating multiple physiologic behaviors. Future studies will focus on identifying the G_i/o_-coupled GPCRs that signal via Gβγ-SNARE implicated in these behavioral and electrophysiological studies, and determining transcriptional signatures and pathologies related to this mechanism.

## METHODS

### Transgenic embryo generation

The SNAP25Δ3 mouse was created with assistance from the Vanderbilt Transgenic Mouse and Embryonic Stem Cell Resource in compliance with protocols approved by the Vanderbilt Institutional Animal Care and Use Committee. Transgenic mouse embryos were generated utilizing the CRISPR/Cas9 system. The protospacer targeting construct was generated via a 24-mer oligo with a forward sequence of5’CACCGCAACAAAGATGCTGGGAAG3’ annealed to 5’AAACTTCCCAGCATCTTTGTTGC 3’. 1 μg of px330 vector (Zhang lab, MIT) was digested in a stoichiometric fashion with Bbs1 (NEB) to a final concentration of 50ng/μL for 1h at 37 C. The oligo was then ligated into the digested product with Quick Ligase (NEB) in a one-pot reaction in which oligo was added to a final concentration of 0.4 μM. and ligated for 4m at 25 C. Constructs were verified via Sanger sequencing. The single-stranded homology donor(IDT, Ultramer) was 126bp in length and spanned the C-terminal final exon of SNAP25 with 48bp of homology in either direction of the site of interest along with the G204* mutation and a HindIII site 3’ of the G204* for the purposes of sequencing. The px330 vector and single-stranded homology donor were co-microinjected into the pronucleus of 587 B6D2 embryos, 447 of which were implanted into 40 B6D2 dams. 32 pups were obtained, two of which contained the G204* mutation in germline cells as measured by PCR analysis of genomic DNA. To verify that the inserted transcript was correct, PCR products were then excised and ligated into pCR2.1TOPO, which was then subjected to Sanger sequencing using M13 and T7 primers. No changes other than the addition of the G204* and HindIII site were observed.

### Mouse breeding and genotyping

Littermate cohorts in this study carry a mixed genetic background of 75% C57BL/6J and 25% DBA2. The presence of the G204* mutation in the SNAP25Δ3 mouse was verified via PCR and Sanger sequencing. Genomic DNA was extracted from the fecal epithelium (Zymo Research) of weaned animals, and homozyogous SNAP25Δ3 animals were differentiated from WT littermates using two separate PCR reactions. Genomic DNA was amplified using a common reverse primer GGATTGTGGCAGTAGCTCG with diagnostic forward primers: the SNAP25Δ3-specific forward primer had the sequence 5’ GATGCTGTAAGCTTAGTGG 3’, while the WT-specific forward primer had the sequence 5’ GCAACAAAGATGCTGGGAAGTGG 3’. Amplicons generated are 467 bp and 459 bp respectively for SNAP25Δ3 and WT mice and were analyzed via agarose gel electrophoresis.

### Plasmids-

The open reading frames for syt I 96-421 was subcloned into the glutathione-s-transferase (GST) fusion vector, pGEX6p-1^91^, (GE Healthcare, Chalfont St. Giles, Buckinghamshire, UK) for expression in *E.coli*. t-SNARE complexes consisting of SNAP25 and syntaxin1A were purified from the dual-expression vector pRSF-Duet1 with a subcloned N-terminal-His tag on SNAP25^92^. C-terminally His-tagged synaptobrevin was produced from the plasmid pTW2^48^. The sequence corresponding to a gRNA specific to the final exon of SNAP25 was cloned into pX330-U6-Chimeric_BB-CBh-hSpCas9^93^, which was a gift from Feng Zhang (Addgene plasmid # 42230). t-SNAREs containing SNAP25Δ3 were produced via point mutagenesis of the WT SNAP25 sequence in pRSF-Duet1 using the method of overlapping primers. pCR2.1TOPO was obtained from Invitrogen.

### Immunoblotting

For the immunoblot analysis, SNAP25 (Santa Cruz, sc-376713, 1:500), SNAP23 (Abcam, ab3340,1:500), cysteine string protein (Abcam, ab90499, 1:1000), tomosyn (Santa Cruz, Sc-136105, 1:1000), Hsc70 (Abcam, ab154415, 1:1000), Munc13-1 (Synaptic Systems, 126102, 1:1000), munc18-1 (Abcam, ab3451, 1:3000), synaptotagmin7 (Synaptic Systems, 105173, 1:1000), VAMP2 (Synaptic Systems, 104211, 1:1000), synaptotagmin1 (Synaptic Systems, 105-011, 1:1,000), complexin 1/2 (Synaptic Systems 122002, 1:500), syntaxin-1 (Santa Cruz, sc-12736, 1:2,000), and β-actin (Abcam, ab8227 1:5000) were used. HRP-conjugated secondary antibodies were obtained from Perkin-Elmer and used at the following dilutions: goat anti-mouse (1:10,000), and goat anti-rabbit (1:10,000). Images were analyzed for densitometry using ImageJ (available from http://rsbweb.nih.gov/ij/index.html). All statistical tests were performed using GraphPad Prism v.4.0 for Windows, (GraphPad Software, La Jolla, California, USA, www.graphpad.com).

### Synaptosome preparation, fractionation, and lysate protocol

Crude synaptosomes were prepared, fractionated, and lysed from mouse brain tissue, as described previously^94^. Various synaptic proteins were detected in lysate of whole crude synaptosomes and presynaptic fractions.

### Protein purification and labeling

Recombinant bacterially expressed syntaxin1A and 6xHisSNAP25 (both WT and SNAP25Δ3 mutants) were expressed in tandem and purified from *E.coli* strain BL21, while 6xHisVAMP2 was purified separately. GST-syt I 96-421 was purified separately on glutathione-agarose beads. All four were purified according to previously published methods^27,47,48^. Gβ_1_γ_1_ was purified from bovine retina as described previously^95^. Purified syt I C2AB was buffer exchanged into 25 mM HEPES pH 7.4, 150 mM NaCl, 1 mM TCEP, and 10% glycerol and labeled at a 20-fold excess with Alexa Fluor 488-C5-maleimide at RT for 2H before excess probe was removed with an Amicon centrifugal filter with a molecular weight cutoff of 10,000.

### Preparation of liposomes for fusion and TIRF assays

Small unilamellar liposomes containing t-SNARE complexes or VAMP2 were made as described previously^27,47,48^. A volume of 55% POPC (1-Palmitoyl-2-Oleoyl-sn-glycero-3-phosphocholine), 15% DOPS (1,2-dioleoyl-*sn*-glycero-3-phospho-L-serine (sodium salt) and 30% POPE (1-Palmitoyl-2-Oleoyl-sn-glycero-3-phosphoethanolamine) in chloroform that would be equal to 15 mM of lipids in 100 μl were dried to a lipid film in a glass vial under argon, followed by vacuum removal of residual chloroform. Liposomes containing VAMP2 included 1.5% 1.5% N-(7-nitro-2-1,3-benzoxadiazol-4-yl)-1,2-dipalmitoyl phosphatidylethanolamine (NBD-PE) and 1.5% N-(lissamine rhodamine B sulfonyl)-1,2-dipalmitoyl phosphatidylethanolamine (Rhodamine-PE), along with 55% POPC/15% DOPS/27% POPE. 0.4 mg of t-SNARE dimer or 95uL of VAMP2 was added to each tube of lipids in tandem with elution buffer 25 mM HEPES-KOH; pH 7.8, 400 mM KCl, 500 mM imidazole, 10% glycerol, 5 mM 2-mercaptoethanol, 1% n-octylglucoside) to a final volume of 500 μl and subjected to mild agitation until the lipid film was fully dissolved. Liposomes were then formed via the addition of 2 volumes of reconstitution buffer (25 mM HEPES-KOH, pH 7.8; 100 mM KCl; 1mM DTT; 10% glycerol;) in a dropwise manner, followed by additional mild agitation for 10 minutes. The solution was then dialyzed (10,000 molecular weight cutoff) twice for six hours in 4L of reconstitution buffer to remove residual detergent. Solutions were mixed with equivalent volumes of 80% iohexol (Accudenz, Accurate Chemical Co.) and were purified on a 0%/30%/40% iohexol gradient in a Beckman SW-55 swinging bucket rotor. Liposomes were harvested from the 0-30% interface and flash-frozen at −80°C. Lipid concentrations and recovery rates were obtained using the Beer-Lambert law with NBD-PE absorbance at 460nm from v-SNARE liposomes containing NBD-PE that were maximally dequenched via the addition of docdeylmaltoside to 0.5%. SNARE protein concentrations in VAMP2 and t-SNARE liposomes were determined via Coomassie Brilliant Blue R-250 staining of SDS-PAGE gels containing a standard curve of bovine serum albumin (Thermo Scientific) followed by densitometric analysis of pixel intensity of syntaxin1A bands utilizing the Fiji distribution of ImageJ software^96,97^. SNARE copy number in liposomes was determined according to previously published methods^47^.

### In membrane TIRF imaging

Lipid bilayers were prepared from liposomes containing 55% PC/15%PE/29%PS in addition to 1% DiD (3,3′-Dioctadecyloxacarbocyanine perchlorate) with or without t-SNARE complexes containing WT or mutant SNAP25. TIRF imaging studies were conducted according to previously published methods^27^. A shallow volume (to ∼150 nm above the cover slip) was illuminated using TIRF on a custom microscope utilizing a laser launch with 488 nm solid state and 633 nm HeNe lasers whose output intensities were controlled using a Acoutic-Optical-Transmission Filter (AOTF, Prairie Technologies). 488 nm laser intensities at the microscope input used to illuminate fluorescently tagged proteins were 2 to 5 mW. This beam was offset and focused on the objective backplane (Olympus PlanApoN 60x 1.45 TIRFM) using a modified TIRF laser launch (Olympus IX2). Coverslips were 0.17 mm borosilicate glass. After laying down lipids in a 5 mM CaCl_2_-containing solution, a superfusate of KCl 150 mM and HEPES 5mM washed the lipid. Then AF-sytI with 100 µM free Ca^2+^ was applied in solution over the cover-slip-supported lipid bilayer. The 633 nm excitation was used to focus the TIRF illumination on the bilayer. Simultaneously, the intensity and polarization of 535nm fluorescence emitted from Alexa Fluor 488-labeled syt1 C2AB (AF-syt1) was measured by capturing P and S polarized emission across a polarizing cube beamsplitter^27^. The focus and setting of the TIRF field used a camera in the inverted configuration light path. Emission was detected and quantified through a second water immersion lens to minimize loss of polarity in emission, with excitation light polarity in plane with one detector and orthogonal to another after a polarizing beam splitter.

### Behavioral Studies

### Subjects

All studies were conducted in adult male wildtype and SNAP25Δ3 homozygote mice that were breed in house as described above and group-housed unless specified under a 12/12 h light-dark cycle with water available *ad libitum*. All behavioral studies were evaluated in cohorts of age-matched, littermate controlled wildtype and SNAP25Δ3 homozygote mice. All animal experiments were approved by the Vanderbilt University Animal Care and Use Committee and experimental procedures conformed to guidelines established by the National Research Council *Guide for the Care and Use of Laboratory*

### Animals

All efforts were made to minimize animal suffering and the number of animals used. *Modified Irwin Neurological Test Battery and Body Weights.* This test battery evaluated changes in 25 different autonomic and/or somatomotor nervous system endpoints using the Modified Irwin Neurological Test Battery^50^ in age-matched littermate male SNAP25Δ3 homozygotes (n= 21) or wild-type (n= 17) of 14-15 weeks of age. All data were manually collected by an experimenter blinded to genotype. Data were represented as the aggregate mean scores for all of the autonomic or somatomotor nervous system endpoints of the Irwin test battery per genotype and were analyzed by Student’s t-test using GraphPad Prism. Differences were evaluated using a rating scale from 0-2 with 0=no effect, 1=modest effects, 2=robust effect. The aggregrate mean scores were extrapolated by calculating the sum of each Irwin parameter for all mice tested. Then the mean of all of the autonomic or somatomotor parameters were calculated. Data are represented as means ± SEM and were analyzed by Student’s two-tailed t-test using GraphPad Prism. In addition, total body weight in grams (g) was measured between 3-60 weeks of age for WT and SNAP25Δ3 homozygote mice. Data are presented as means ± SEM and were analyzed by two-way ANOVA using GraphPad Prism.

### Stress induced hyperthermia assay

Singly-housed male SNAP25Δ3 homozygotes or wild-type littermate controls were allowed to acclimate to the testing room for 60 minutes. Baseline temperature was taken by inserting a BAT-12 Microprobe Thermometer, dipped into mineral oil, 2 cm into the rectum for each mouse for 20 s to obtain core body temperature (*T*_1_). After a 15-min interval, core body temperature for each mouse was measured a second time (*T*_2_). Stress-induced hyperthermia was calculated as the change in core body temperature between the first and second temperature readings (Δ*T* = *T*_2_ - *T*_1_) based on previously published methods^61^ Data are presented as means ± SEM and were analyzed by Student’s two-tailed t-test using GraphPad Prism.

### Locomotor Activity Assay

Open field activity was tested in age-matched 10-12 week old male WT (n=15) and homozygote SNAP25Δ3 animals (n=14) using an open field system (OFA-510, MedAssociates, St. Albans, VT) with three 16 × 16 arrays of infrared photobeams as previously described^98^. Total number of photobeam breaks were collected over 60 min under light or infrared lighting conditions. Data were calculated as either the mean distance traveled (cm) ± SE for the number of photobeam breaks/5 min bin/genotype, the total distance traveled (mean horizontal and vertical beam breaks/30 mins) or the total rearing behavior (mean vertical beam breaks/30 mins. Data are presented as means ± SEM and were analyzed by two-way ANOVA (mean distance traveled) and Student’s two-tailed t-test (total distance traveled and rearing) using GraphPad Prism.

### Tail-Flick Test

Age-matched littermate male SNAP25Δ3 homozygotes or WT controls were assessed in the tail-flick assay, a preclinical model of acute spinal-mediated thermal nociception^56^. All mice had their tails individually immersed in a water bath maintained at 55°C, and the latency to removal in sec was measured. If an animal did not flick its tail within 10 s, it was removed and assigned a response time of 10 s. Data are presented as means ± SEM and were analyzed by Student’s two-tailed t-test using GraphPad Prism.

### Hot Plate Test

Age-matched littermate male SNAP25Δ3 homozygotes or WT controls were assessed in the hot plate assay, a preclinical model of acute supraspinal-mediated thermal nociception^87^. All mice were individually placed on a hot plate maintained at 50-55°C and the latency to lick the front or hind paws was recorded for each mouse. Animals not responding within 30 s were removed and assigned a score of 30 s. Data are presented as means ± SEM and were analyzed by Student’s two-tailed t-test using GraphPad Prism.

### Rotarod assay

Age-matched 10-12 week old male WT (n=15) and homozygote SNAP25Δ3 animals (n=14) were assessed for their ability to maintain balance upon a rotating cylinder undergoing constant acceleration from 4 to 40 RPM^99^. The cylinder was 3cm in diameter and each animal was confined to approximately 6cm of cylinder length via the use of Plexiglas dividers. Cylinders were suspended 25cm above a lever that actuates the timer, resulting in a stoppage of the timer to permit tabulation of the latency required for the animal to fall off the cylinder. The maximum length for each trial was 300 sec, at which point a maximum latency value of 300 was tabulated. Trials were conducted for three consecutive days. Data for each day were represented as the mean latency to drop from the rotarod in sec ± SE and analyzed using Student’s two-tailed t-test for comparisons by day using GraphPad Prism.

### TreadScan^®^ Gait Analysis System

Abnormalities in gait in male SNAP25Δ3 homozygotes or wild-type littermate controls were assessed with the TreadScan^®^ system (CleverSys, Inc., Reston, VA)^53^. Animals were placed on a transparent treadmill and made to run at treadmill speeds (mm/sec) of 120 or 160, while video footage of the animal’s paws was recorded. The animals were tested starting from the lowest to highest speed with a 20 minute interval between each test. Video footage was analyzed for the position of each paw utilizing the Treadscan software suite to assess any abnormalities in gait. Data were represented as the mean ± SE for the following parameters: brake duration (msec), stance duration (msec), gait angle (^o^), track width (mm), stride frequency (Hz), stride length (mm), % stance, stride duration (msec), swing duration (msec), propel duration (msec), and run speed (mm/s) and analyzed using Student’s two-tailed t-test using GraphPad Prism.

### Light-dark exploration

Anxiety responses were assessed in age-matched WT or SNAP25Δ3 homozygote male mice in a plastic cubic light-dark chamber 20 cm in length^54^. A black opaque Plexiglas insert was utilized to selectively create darkness in half of the chamber. Animals were placed individually in each chamber for 5 minutes, with animal movement being recorded by the breaking of photobeams emitted by infrared photocells in each chamber. Time spent in the light side of the chamber (sec), number of transitions made by each mouse between the light and dark sides of the chamber, and distance traveled in the light and the dark sides of the testing chamber (cm) were recorded. Data are represented as means ± SEM and were analyzed by Student’s two-tailed t-test using GraphPad Prism.

### Forced Swim Task

Age-matched littermates 16-17-week old WT (n= 14) or SNAP25Δ3(n=17) homozygote male mice were tested in forced swim paradigm^55^. The depth of the water was such that the mice tail does not touch the bottom, but also prevents them from escaping the apparatus. For testing, each mouse was placed in the cylinder for 6 mins, and the latency to immobility and the immobility time (i.e. the time during which mice made only the small movements necessary to keep their heads above water) was scored. Only the data scored during the last 4 min were analyzed. Data are represented as means ± SEM and were analyzed by Student’s two-tailed t-test using GraphPad Prism.

### Morris Water Maze

Mice were trained to locate a hidden platform in a standard fixed platform memory acquisition task, in which the platform remained in a constant position, see methods^96^, ^97^. This acquisition phase lasted for six sessions, each of which consisted of four trials separated by approximately 10 min. Four points along the perimeter of the maze, served as starting points where the mice were released, facing the wall of the tank, at the beginning of each trial (the order of the starting points were determined randomly, except that each starting point was used only once each session). After a mouse located the platform, it was allowed to remain there for 30 s before being removed from the tank. If a mouse failed to locate the platform within 60 s, it was manually guided to it and again allowed to remain on the platform for 30 s. Data are represented as means ± SEM and were analyzed by Two-way ANOVA using GraphPad Prism.

### BNST Electrophysiological analyses

Adult mice (>8 weeks) were single housed in the institutional vivarium prior to experiments. Food and water were available ad libitum. All procedures were approved by the Animal Care and Use Committee at Vanderbilt University. Mice were transported from the animal colony to the laboratory and allowed to acclimate in a sound-attenuation chamber for >1 hour before the experiment. They were then anesthetized with isoflurane until unresponsive to foot pinch. The mice were transcardially perfused with ice-cold sucrose-substituted artificial cerebrospinal fluid (ACSF) (in mM: 194 sucrose, 20 NaCl, 4.4 KCl, 2 CaCl_2_, 1 MgCl_2_, 1.2 NaH_2_PO_4_, 10.0 glucose, and 26.0 NaHCO_3_) saturated with 95% O_2_/5% CO_2_. They were then decapitated and the brain was quickly removed and placed in ice-cold sucrose ACSF. Slices 300 μm thick containing the bed nucleus of the stria terminalis (bregma +0.26 to +0.14 mm) were prepared using a Tissue Slicer (Leica). After dissection, slices were transferred to a holding chamber containing heated (∼29°C) oxygenated (95% O_2_/5% CO_2_) ACSF (in mM: 124 NaCl, 4.4 KCl, 2 CaCl_2_, 1.2 MgSO_4_, 1 NaH_2_PO_4_, 10.0 glucose, 26.0 NaHCO_3_; pH 7.2-7.4; 295-305 mOsm). Recording electrodes (0.5-2 MΩ) were pulled on a Flaming-Brown Micropipette Puller (Sutter Instruments) using thin-walled borosilicate glass capillaries. Excitatory field potentials were evoked by local fiber stimulation with bipolar nichrome electrodes. Electrical stimulation (1-15 V with 5 ms duration) was applied at 0.0167 Hz. Recording electrodes were filled with ACSF and all experiments were done in the presence of 25 μM picrotoxin to isolate fast excitatory transmission. Signals were acquired via a Multiclamp 700B amplifier (Axon Instruments), and digitized and analyzed via pClamp 10.2 software (Axon Instruments). The fiber volley potential (N1) was monitored continuously throughout the duration of the experiment. Experiments in which N1 changed by 20% in either direction were not included in subsequent analysis. Experiments were analyzed by measuring peak amplitudes of the N1 and N2 (synaptic potential) relative to the amplitude with no stimulation. This measure was then normalized to the last ten minutes of the baseline period (10-20 minutes). Welch’s t-test was used to compare the average amplitude over the last 20 minutes of the experiment relative to baseline across the two genotypes.

### Subiculum Electrophysiological analyses

All procedures were approved by the Animal Care and Use Committee at Vanderbilt University and at the University of Illinois at Chicago. Male and female adult mice (>8 weeks) were obtained from the institutional vivarium prior to experiments, where food and water were available ad libitum. Mice were transported from the animal colony to the laboratory and allowed to acclimate for >1 hour before the experiment. They were then anesthetized with isoflurane until unresponsive to foot pinch. The mice were decapitated and the brain was quickly removed and placed in ice-cold sucrose-substituted artificial cerebrospinal fluid (ACSF) (in M: 11 D-Glucose, 234 sucrose, 2.5 KCl, 1.25 NaH_2_PO_4_, 10 MgSO_4_, and 26 NaHCO_3_; in mM: 0.5 CaCl_2_,) saturated with 95% O_2_/5% CO_2_. Slices 300 μm thick containing the hippocampus and subiculum were prepared using a Tissue Slicer (Leica VT1200S). After dissection, slices were transferred to a holding chamber containing oxygenated (95% O_2_/5% CO_2_) ACSF (in mM: 123 NaCl, 26 NaHCO_3_, 3 KCl, 1.25 NaH_2_PO_4_, 11 D-Glucose; 2 CaCl_2_, 1 MgCl_2_; pH 7.2-7.4; 295-305 mOsm) for 45 minutes in a heated (∼30°C) water bath then moved to room temperature for storage. Recording electrodes (3-5 MΩ filled with ACSF) were pulled on a Flaming-Brown Micropipette Puller (Sutter Instruments) using standard-walled borosilicate glass capillaries. Excitatory field potentials were evoked by local fiber stimulation with bipolar twisted insulated NiChrome electrodes in an interface chamber (Warner, BSC-BUW) heated to ∼30°C. Electrical stimulation (200µs at 30 to 60 µA range) was applied every 30 seconds. All experiments were done in the presence of 10 μM bicuculline to black GABA_A_ receptors and 50 µM D-2 amino-5-phosphonopentanoate (D-AP5) to block NMDA receptors to isolate AMPAR-mediated EPSPs. Signals were acquired via an Axoclamp 2B amplifier (Axon Instruments) amplified 100 x and band pass filtered (0.1 to 5KHz), and digitized and analyzed via AxographX version 1.6.5 software. Experiments were analyzed by measuring the initial slope of the excitatory postsynaptic potential and then normalized to the last ten minutes of the baseline period (20 minutes). Student’s t-test was used to compare the average slope over the last 10 minutes of each dose of CP93129 (0, 50, 100, 200, 400, 800, and 100 nM) or baclofen (0, 0.5, 1, 2, 5, and 10 μM) concentrations relative to baseline across the wild-type and SNAP25Δ3 genotypes.

## Supplementary Tables and Figures

**Figure S1. Table of neurobehavioral parameters screened in Modified Irwin Neurological Test Battery. A1**. Aggregate scores of autonomic component of Irwin neurobehavioral screen conducted in 14-15 week old age-matched WT and SNAP25Δ3 homozygote male mice. (n=17-21; ***p<0.001). **A2**. Aggregate scores of somatomotor component of Irwin neurobehavioral screen conducted in male WT and SNAP25Δ3 homozygotes (n=17-21, ****p<0.0001). **B**. The effects of WT (n=17) were compared to littermate SNAP25?3 homozygotes (n=21) on the autonomic nervous system and somatomotor function. The mean scores for each parameter are classified as follows:-,=no effect; (+), 0.01-0.25; +, 0.251-0.50; ++, 0.51-1.0; +++, 1.01-1.50; ++++, 1.51-2.0;**p<0.01 vs. vehicle (Student’s two-tailed t-test).

**Figure S2. SNAP25Δ3 homozygotes do not display abnormalities in unstressed motor function. A**. Plot of distance traveled in five-minute intervals in a dark open field for littermate WT and SNAP25Δ3 homozygotes 16 weeks of age. Total distance traveled and number of rearing movements are plotted below. No significant differences by genotype were seen in this paradigm as measured by two-way ANOVA.

**Figure S3. Additional gait parameters studied in the SNAP25Δ3 homozygote**. A-I: Tabulated parameters for each paw as measured by TreadScan^®^ gait analysis in which the paws of a running male WT or SNAP25Δ3 homozygote are imaged by a camera located beneath a transparent treadmill moving at a speed of 120 or 160 mm/s. Parameters, such as gait angle, track width, stride frequency, stride length, % stance, stride duration, swing duration, propel duration and run speed exhibited no or very modest differences between WT and SNAP25Δ3 homozygotes (*, p<0.05 Student’s two-tailed t-test). Animal cohort sizes were identical to **Figure 4**.).

**Figure S4: SNAP25Δ3 homozygotes do not display elevated anxiety in the absence of external stressors. A-B**. Bar graphs showing time spent by 24-week old littermate male WT (n= 14) or SNAP25Δ3 homozygotes (n=16) in each side of a bipartite chamber in which one half was darkened through the use of a Plexiglass divider. Data were represented as time spent in the light side of the chamber (sec), number of transitions made by each mouse between the light and dark sides of the chamber, and distance traveled in the light and the dark sides of the testing chamber (cm) as denoted by the number of infrared beam breaks within the apparatus in dark or light portion of the testing apparatus. No significant differences were detected in time spent in each side of the chamber (p= 0.5209) or number of transitions made (p= 0.2601).

## ACKNOWLEDGEMENTS

The work in this paper was supported by R01EY010291; R01MH101679 and R01DK109394 to HEH; R01MH 08487 to STA; R01DA042475 and F30DA042501 to DGW. We thank Jennifer Skelton and the Vanderbilt Transgenic Mouse and Embryonic Stem Cell Resource (supported by NIH grants CCSG CA68485 and DRTC DK020593) for assistance in the creation of the SNAP25Δ3 mouse. We thank the Vanderbilt High-throughput Screening Core for providing instrumentation for lipid mixing experiments. We thank John Allison, Ph.D and the Vanderbilt Murine Neurobehavioral Core for providing training and instrumentation for the behavioral studies conducted upon the SNAP25Δ3 mouse. We thank Bob Matthews Ph.D and the Vanderbilt Cell Imaging Shared Resource (supported by NIH grants CA68485, DK20593, DK58404, DK59637, and Ey08126) for training and instrumentation in microscopy. We thank Mark Fulton and Kayla Jo Temple for assistance in liposome production, and the laboratory of Feng Zhang, Ph.D (Massachusetts Institute of Technology) and Addgene for the gift of the px330 plasmid.

## AUTHOR CONTRIBUTIONS

Conceptualization, Z. Z., D.P.M., D.G.W., S.A., C.K.J.,., and H.E.H. Formal Analysis, Z. Z., A. D. T. G., L. J. B., YY. Y., N. A. H., and S. A. Methodology, Z. Z., A.D.T.G., YY.Y., D.G.W., S.A., C.K.J., and H.E.H. Investigation, Z. Z., A.D.T.G., L.J.B., B.P., E.C., N.A.H., M.R.D., YY.Y. and S. A. Writing-Original Draft, Z. Z., A.D.T.G., L.J.B., N.A.H., YY.Y., D.G.W., S.A., C.K.J., and H.E.H. Writing-Review and Editing, Z. Z., A.D.T.G., L.J.B., N.A.H., YY.Y., D.G.W., S.A., C.K.J., and H.E.H. Resources, Z. Z., A.D.T.G., K.H., L.J.B., and D.P.M. Funding Acquisition, H.E.H. Supervision, D.G.W., S.A., C.K.J., and H.E.H.

